# Role of the antiparallel double-stranded filament form of FtsA in activating the *Escherichia coli* divisome

**DOI:** 10.1101/2024.06.24.600433

**Authors:** Abbigale Perkins, Mwidy Sava Mounange-Badimi, William Margolin

## Abstract

The actin-like FtsA protein is essential for function of the cell division machinery, or divisome, in many bacteria including *Escherichia coli*. Previous *in vitro* studies demonstrated that purified wild-type FtsA assembles into closed mini-rings on lipid membranes, but oligomeric variants of FtsA such as FtsA^R286W^ and FtsA^G50E^ can bypass certain divisome defects and form arc and double-stranded (DS) oligomeric states, respectively, which may reflect conversion of an inactive to an active form of FtsA. Yet, it remains unproven which oligomeric forms of FtsA are responsible for assembling and activating the divisome. Here we used an in vivo crosslinking assay for FtsA DS filaments to show that they largely depend on proper divisome assembly and are prevalent at later stages of cell division. We also used a previously reported variant that fails to assemble DS filaments, FtsA^M96E R153D^, to investigate the roles of FtsA oligomeric states in divisome assembly and activation. We show that FtsA^M96E R153D^ cannot form DS filaments *in vivo*, fails to replace native FtsA, and confers a dominant negative phenotype, underscoring the importance of the DS filament stage for FtsA function. Surprisingly, however, activation of the divisome through the *ftsL*\* or *ftsW*\* superfission alleles suppressed the dominant negative phenotype and rescued the functionallity of FtsAM96E R153D. Our results suggest that FtsA DS filaments are needed for divisome activation once it is assembled, but they are not essential for divisome assembly or guiding septum synthesis.

**IMPORTANCE:** Cell division is fundamental for cellular duplication. In simple cells like *Escherichia coli* bacteria, the actin homolog FtsA is essential for cell division and assembles into a variety of protein filaments at the cytoplasmic membrane. These filaments help to tether polymers of the tubulin-like FtsZ to the membrane at early stages of cell division, but also play crucial roles in recruiting other cell division proteins to a complex called the divisome. Once assembled, the *E. coli* divisome subsequently activates synthesis of the division septum that splits the cell in two. One recently discovered oligomeric conformation of FtsA is an antiparallel double stranded filament. Using a combination of in vivo crosslinking and genetics, we provide evidence suggesting that these FtsA double filaments have a crucial role in activating the septum synthesis enzymes.

## INTRODUCTION

Bacterial cytokinesis relies on a dynamic protein nanomachine called the divisome, which is organized in a ring-like complex at mid-cell by the tubulin-like FtsZ protein (1–3). Actin-like FtsA protein plays a crucial role during early FtsZ-ring formation by anchoring FtsZ to the cytoplasmic membrane at the proto-ring stage (Fig. 1) and facilitating the recruitment of other essential divisome proteins (4, 5). Later in cytokinesis, synthesis of the division septum in most bacteria requires the action of the transmembrane proteins FtsW and FtsI, which harbor essential septal glycosyltransferase and transpeptidase activities, respectively, in their periplasmic domains (6–8). FtsWI in turn are recruited to the divisome by the FtsQLB transmembrane protein complex (9–11) and are kept in an inactive state by the periplasmic domains of FtsL and FtsB, collectively called constriction control domains (12). Thus, *E. coli* cell division features two temporally distinct stages (13): first assembly of the divisome machinery, then activation of FtsWI to synthesize the division septum.

**Fig. 1.**
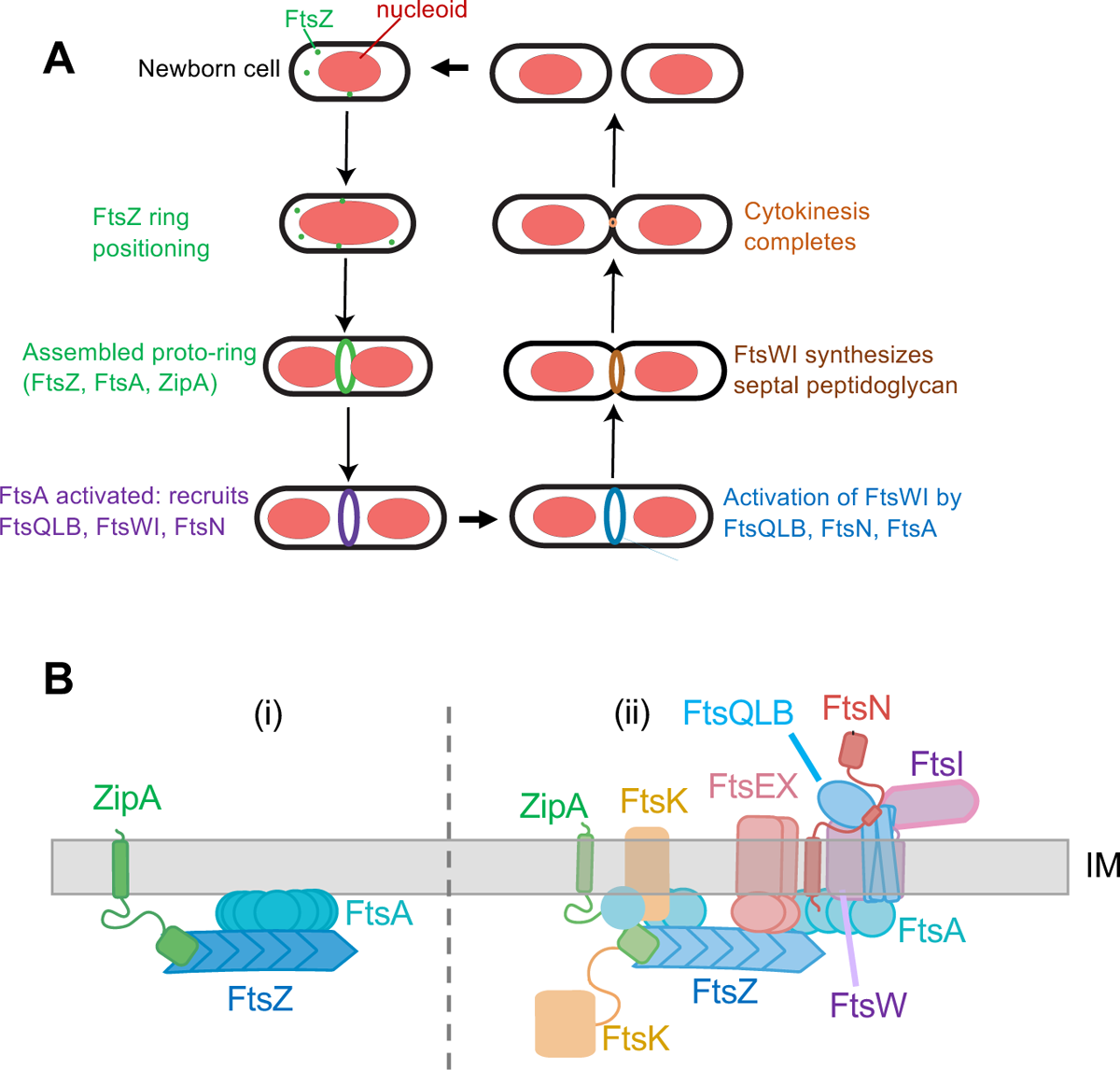
Stages of cell division in *E. coli*. A) Scheme of the cell division cycle, highlighting stages relevant to this study. B) Essential divisome proteins, depicting the known FtsQLBWI complex with FtsN, along with other divisome proteins at the membrane. The proto-ring assembles first (i), followed by the remainder of the divisome (ii).

Significant progress has been made recently in understanding how the *E. coli* divisome is assembled and FtsWI becomes activated. Hypermorphic alleles of FtsA, designated FtsA*, were initially found capable of bypassing the essential ZipA protein as well as other divisome defects (14–16). These FtsA* alleles were found to be deficient in oligomer formation compared with WT FtsA, according to two-hybrid studies (17). Subsequent *in vitro* studies demonstrated that purified FtsA forms closed 12-mer rings on lipid membranes, whereas FtsA* variants instead tended to assemble into arcs and other non-miniring structures, including double stranded (DS) filaments under certain conditions (18, 19). FtsA* also was shown in reconstitution experiments with FtsZ to form more dynamic structures on lipid membranes compared with WT FtsA (20). These biochemical and genetic studies suggested that FtsA oligomeric state regulates a divisome checkpoint, and disruption of FtsA minirings inactivates the checkpoint and allows FtsQLB and FtsWI to be recruited to the divisome (1, 17).

As mentioned above, FtsQLB themselves are key regulators of FtsWI activity. This was initially revealed by hypermorphic alleles of FtsB and FtsL, designated FtsB* and FtsL*, which harbor residue changes in their periplasmic domains that prematurely and constitutively activate FtsWI (12, 21). The FtsQLB complex was later shown to be sufficient to promote FtsWI enzymatic activity *in vitro* (6). In the periplasm, this activation is mainly mediated through a domain in FtsL (AWI) that interacts with FtsI (22). Moreover, hypermorphic variants of FtsW and FtsI, FtsW* and FtsI* have been isolated that mimic the activated state induced by FtsL* and FtsB* (22–24), suggesting that FtsWI activation is the endpoint of these regulatory pathways. Like the original FtsA* allele, these hypermorphic alleles of FtsL, FtsB, FtsW and FtsI promote septum formation prematurely and at shorter cell lengths, resulting in significantly shorter cells (12, 21, 25). Combinations of some of these hypermorphic alleles synergize to form nearly spherical cells (12), presumably because the septum synthesis activity exceeds activity of the elongasome, which is normally dominant and results in the rod shape of *E. coli* and other bacilli (26, 27).

The prevailing model for how FtsA functions in *E. coli* cell division involves several oligomeric state changes, which have been observed *in vitro* but largely only inferred *in vivo* (1, 28). In this model, FtsA initially self-interacts strongly to form minirings on the membrane (19, 29, 30). Upon a signal yet to be elucidated, it is postulated that these FtsA minirings are then disrupted by interactions with the other essential FtsZ membrane anchor protein ZipA (31–33) and perhaps other partners such as the divisome protein FtsX (34) or FtsQ (35) to form curved oligomers (arcs) that, unlike minirings, seem to allow recruitment of downstream divisome proteins, including FtsN.

The cytoplasmic domain of FtsN (FtsN_cyto_), potentially in partnership with other divisome proteins, interacts directly with FtsA (36, 37). Recently it was discovered that high concentrations of purified FtsN_cyto_ can convert FtsA minirings or arcs into DS filaments on lipid membranes (30). These DS filaments are antiparallel, similar to DS filaments formed by another bacterial actin homolog, MreB (30, 38) that guide the peptidoglycan synthesis machinery of the elongasome to maintain the cylindrical shape of the bacterial cell wall (39). Notably, the FtsA^G50E^ variant can form DS filaments on membranes constitutively, independently of FtsN_cyto_ (19, 30). This behavior suggested that FtsA^G50E^, originally isolated as an FtsA*-like allele that suppressed a thermosensitive allele of FtsA (FtsA^S195P^) in *cis* (40), represents another activated oligomeric form of FtsA. As the majority of FtsN is recruited at a late stage of divisome assembly and is important for divisome activation (41, 42), it is attractive to propose that DS filaments reflect an endpoint oligomeric state of FtsA that acts late in septum formation. Because they are curved and oligomerize along the inner membrane surface, these DS filaments were postulated to be crucial for guiding septum synthesis (30). Their antiparallel structure abrogates their ability to treadmill, so Nierhaus et al. hypothesized that DS filaments guide septum synthesis by acting as rudders to steer septal wall synthesis driven by FtsWI, analogous to the rudder mechanism proposed to confer the spatial regulation of the elongasome by MreB DS filaments (30, 39). In the present study, we ask whether these DS filaments of FtsA are required for physically guiding septum synthesis, as proposed. We use site-specific cysteine crosslinking to show that FtsA DS filaments form later in cell division and largely depend on divisome assembly, consistent with the idea that DS filaments are mainly assembled close to the divisome activation stage. We show that the DS filament-forming FtsA^G50E^ variant does not bypass ZipA very efficiently compared with FtsA^R286W^, consistent with the idea that DS filaments act at the later activation stage. Moreover, we show that a variant of FtsA unable to form DS filaments is not only non-functional in cell division but also exhibits a dominant negative phenotype, which cannot be alleviated by incorporating FtsA*-like mutations. Surprisingly, however, this variant can support normal divisome assembly and cell division if the divisome is activated by *ftsL*\* or *ftsW*\* hypermorphic alleles. This leads us to conclude that although DS filaments of FtsA are not essential for guiding septum synthesis, they play a key role in the activation of the divisome.

## RESULTS

### Q155C can be used to detect the double stranded filament form of FtsA by *in vivo* crosslinking and does not affect normal FtsA function

Previously, it was shown that expression of the FtsA^Q155C^ variant could detect DS filaments of FtsA in *E. coli* cells by site-specific cysteine crosslinking with another Q155C residue on the antiparallel FtsA filament (30). As the distance between the Q155C residues in each FtsA filament is 12.6 Å in the atomic structure (Fig. 2A), thiol crosslinkers with sufficiently long linker arms, such as BMH (13-16 Å) or BMB (11 Å), could efficiently capture the interaction and resulted in a single prominent crosslinked species on immunoblots from SDS-PAGE (30). Nierhaus et al. showed that this *ftsA*^Q155C^ variant, fused to an internal 3X HA tag for immunodetection, was able to replace the WT *ftsA* at the native locus of *E. coli*; these cells divided normally, indicating that the Q155C residue change did not detectably affect FtsA function (30). We confirmed that when expressed from a plasmid, FtsA^Q155C^ with an N-terminal FLAG tag (but lacking the 3X HA tag used previously) can complement an *ftsA* null allele and confer normal cell division (data not shown and see below). We then used this site-specific crosslinked band as a direct measure of the prevalence of FtsA DS filaments in cells under various conditions.

**Fig. 2.**
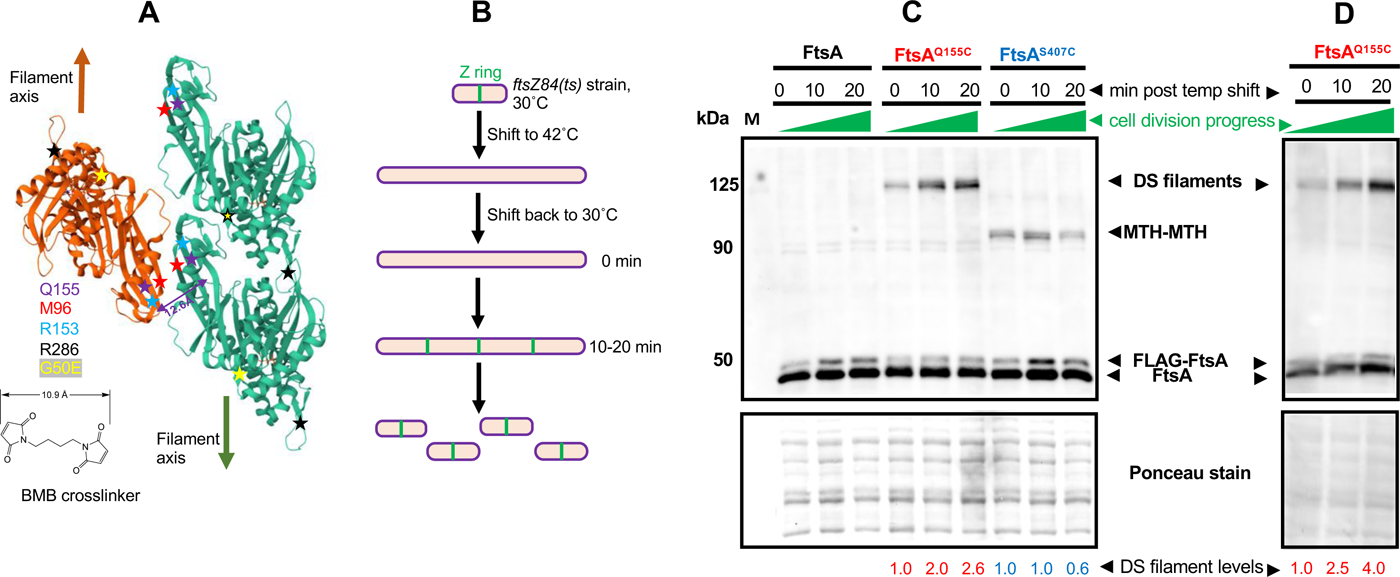
*In vivo* crosslinking detects an increase in FtsA DS filaments as cell division progresses. (A) Diagram of three FtsA subunits within an antiparallel DS filament (Nierhaus et al. 2022) showing two green subunits in one filament and one orange subunit in the opposing filament) and positions of residues described in this study, including the closely apposed Q155 residues that were replaced with cysteines and used for cysteine crosslinking by BMB or BMH crosslinkers. The structure was derived from PDB 7Q6F. The BMB crosslinker structure is depicted; BMH crosslinker is similar but spans 13 Å. S407 is part of the C-terminal membrane-binding amphipathic helix and was not present in the crystal structure. (B) Scheme used to synchronize *E. coli* cell division, using the *ftsZ84*(ts) allele. *E. coli ftsZ84*(ts) cells (WM1125) were transformed with pDSW210FLAG-FtsA (negative crosslinking control), -FtsA^Q155C^ (detects DS filaments by crosslinking between cysteines) or -FtsA^S407C^ (detects general FtsA-FtsA intersubunit interactions through MTH-MTH interactions by cysteine crosslinking). After growth at 30°C to early logarithmic phase, cells were shifted to 42°C to disrupt Z rings, then shifted back to 30°C to rapidly reassemble Z rings and restart the cell division cycle. C) Blot of samples at various time points after shifting back to 30°C, separated by SDS-PAGE and probed with anti-FtsA antibody. DS filament crosslinks run at ∼125 kDa, whereas MTH-MTH crosslinks run at the FtsA dimer size of ∼90 kDa. Note that the lower levels of FLAG-FtsA compared with FtsA is in part a result of cleavage of the FLAG tag from the plasmid-expressed version. A portion of the Ponceau S-stained blot is shown below the immunoblot, along with relative intensities of the crosslinked band normalized to the zero minute time point. D) Blot from an independent experiment using cells from strain WM7547 carrying pDSW210FLAG-FtsA^Q155C^ introduced into the *ftsA° ftsZ84*(ts) strain WM7412, with the same protocol as in (B), showing a similar increase in crosslinking over time. A portion of the Ponceau S-stained blot is shown below the immunoblot, along with relative intensities of the crosslinked band normalized to the zero minute time point for each Q155C or S407C time point experiment shown.

### DS filaments of FtsA increase at later stages of division

The prediction inferred from the ability of FtsN_cyto_ to convert FtsA minirings into DS filaments is that DS filaments appear later in the septation process. To test this idea, we measured the prevalence of DS filaments during a synchronized cell division time course. We achieved transient cell division synchrony by using the well-known *ftsZ84*(ts) allele, which rapidly inactivates Z rings upon temperature shift to 42°C and rapidly restores them upon shift back to 30°C (25). As a result of this restoration, divisomes are rebuilt rapidly, and nondividing cells growing in rich medium such as LB generally begin dividing again 20-30 minutes after the downshift (Fig. 2B and data not shown). We therefore used these reactivation conditions to determine whether FtsA DS filaments assembled late during the cell division cycle as hypothesized.

Cells with the *ftsZ84*(ts) allele and carrying pDSW210-FLAG-FtsA^Q155C^ were shifted to 42°C to disassemble divisomes and then harvested for cysteine crosslinking 0, 10, and 20 minutes after returning to 30°C. Protein samples from these time points were separated by SDS-PAGE, followed by immunoblotting with anti-FtsA antibody to detect FtsA. We found that the FtsA^Q155C^ crosslinked band, initially nearly undetectable, increased several-fold in intensity over time at 30°C, consistent with the assembly of most DS filaments at late stages of septation (Fig. 2C,D). This band consistently migrated at ∼125 kDa with SDS-PAGE, which is larger than the predicted FtsA dimer size of ∼90 kDa; however, unexpectedly slow migration of crosslinked bands with SDS-PAGE is often observed with other *in vivo* crosslinking assays (43) and was previously observed specifically with FtsA^Q155C^ (30).

To independently assess FtsA-FtsA interactions not directly involved in DS filaments, we constructed FtsA^S407C^, with a cysteine substitution in the amphipathic membrane targeting helix (MTH) FtsA uses for binding to the membrane (4). Examination of high resolution tomograms of negatively stained minirings on lipids suggests that these MTHs could potentially form an internal ring inside the 20 nm-diameter miniring, potentially placing them in close contact with each other (18). We also hypothesized that at least a subset of MTHs should similarly interact with each other at the membrane surface when FtsA is assembled in other oligomeric structures. Indeed, the immunoblots indicated that FtsA^S407C^ can efficiently crosslink with FtsA^407C^ residues on other FtsA subunits when BMB or BMH crosslinkers are used, yielding crosslinked bands at ∼90 kDa, the expected size of an FtsA dimer (Fig. 2C). As with FtsA^Q155C^, FtsA^S407C^ complemented an *ftsA* null mutant and allowed normal cell division (data not shown), indicating that the serine to cysteine change did not appreciably disrupt the amphipathic nature of the helix. Notably, whereas FtsA^Q155C^ crosslinking increased significantly upon resumption of cell division, FtsA^S407C^ crosslinking remained relatively constant over time, indicating that the observed increase in FtsA^Q155C^ crosslinking arises from specific FtsA-FtsA lateral interactions at the DS filament interface and not from an overall increase in FtsA-FtsA interactions as cell division progresses. We also tested another crosslinking residue, D123C, which like Q155C detects DS filaments *in vivo* but with a much closer molecular interaction (∼3.6 Å) (30). As a result, this interaction cannot be detected by BMB or BMH but can be detected by the shorter ∼5 Å crosslinker dBBR (30). As with Q155C, we saw a 2-3-fold increase in crosslinking at D123C after resumption of cell division in the *ftsZ84* strain, although the efficiency of crosslinking was lower (Fig. S1).

### FtsA DS filaments depend on normal divisome assembly

To provide more evidence that DS filaments of FtsA form at a later stage of the cell division cycle, we asked whether DS filament formation would be hindered by disrupting divisome assembly. To assess this, we introduced pDSW210-FLAG-FtsA^Q155C^ into strains with *zipA1*(ts) or *ftsQ1*(ts) alleles, as both fail to recruit downstream divisome proteins at the nonpermissive temperature of 42°C (44–46). We then compared the intensity of crosslinked bands on immunoblots at the permissive or nonpermissive temperatures. Additionally, we included a strain carrying the pDSW210-FLAG-FtsA^Q155C^ with an *ftsA* null allele in the chromosome as a control to account for the effects of thermal induction.

When grown at 30°C, all three strains displayed the expected prominent crosslinked bands at ∼125 kDa (Fig. 3). However, after shifting to 42°C for 1 h, the intensities of the crosslinked bands in the *zipA1* and *ftsQ1* mutants decreased 7.5-fold and 3-fold, whereas the intensities of the bands in the complemented *ftsA* mutant remained nearly the same. As essentially all *zipA1* and *ftsQ1* cells were highly filamentous at this time point at 42°C (data not shown), it is possible that the weak but detectable crosslinked protein bands represent DS filaments that formed before the thermal shift, which failed to disassemble.

**Fig. 3.**
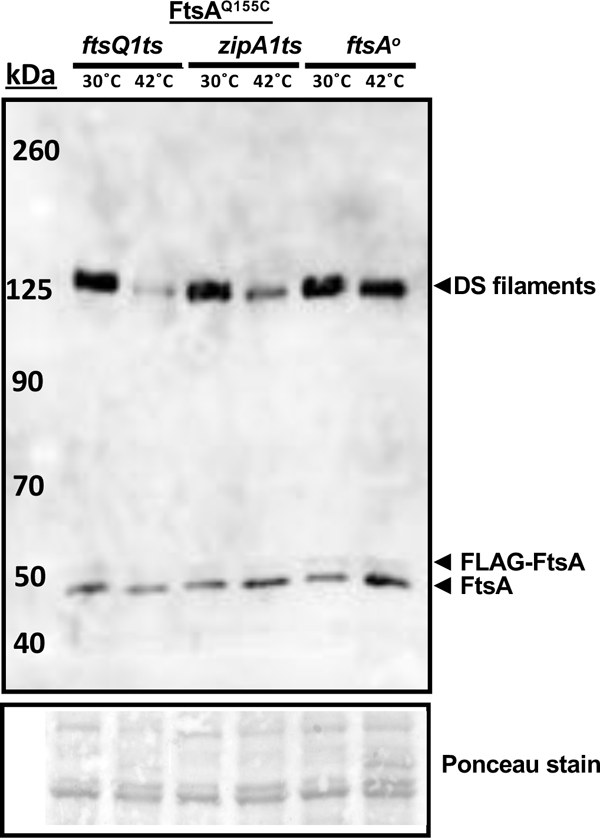
DS filament formation depends on divisome assembly. Derivatives of plasmid pDSW210-Flag-FtsA^Q155C^ were introduced into *ftsQ1ts* (WM7148), *zipA1ts* (WM7149) or combined with an *ftsA°* allele (WM7150), grown at 30°C until early logarithmic phase, then split into 30°C or 42°C cultures for an additional hour prior to cysteine crosslinking before SDS-PAGE and immunoblotting with anti-FtsA. A portion of the Ponceau S-stained blot is shown below the immunoblot.

Overall, these data suggest that formation of FtsA DS filaments largely depends on assembly of the divisome, at least up to the FtsQ-dependent step. As ZipA is important for recruiting all divisome proteins downstream of FtsA and FtsZ and is postulated to be involved in switching FtsA minirings to arcs and/or DS filaments, our results are consistent with this model. They are also consistent with the increase in DS filaments at later stages of septation, as shown above.

### Residue changes that promote or block FtsA DS filament formation *in vivo*

We hypothesized that the cysteine crosslinking assay should be able to detect higher levels of crosslinking in cells expressing FtsA^G50E^ compared to WT FtsA, assuming FtsA^G50E^ forms DS filaments constitutively *in vivo*. We therefore compared the crosslinked bands on immunoblots, and indeed FtsA^G50E^ ^Q155C^ reproducibly gave rise to more intense crosslinked protein bands at ∼125 kDa compared with FtsA^Q155C^ alone (Fig. 4, Fig. S2). As a control, we tested whether the presence of G50E affected the MTH-MTH interaction probed by crosslinking at S407C, but there was no increase in band intensity at ∼90 kDa (Fig. 4). This provides additional evidence, already suggested by the time course of crosslinking depicted in Fig. 2, that the intensity of crosslinked protein bands from FtsA^Q155C^ can be used to assess the prevalence of DS filament formation in the cell population.

**Fig. 4.**
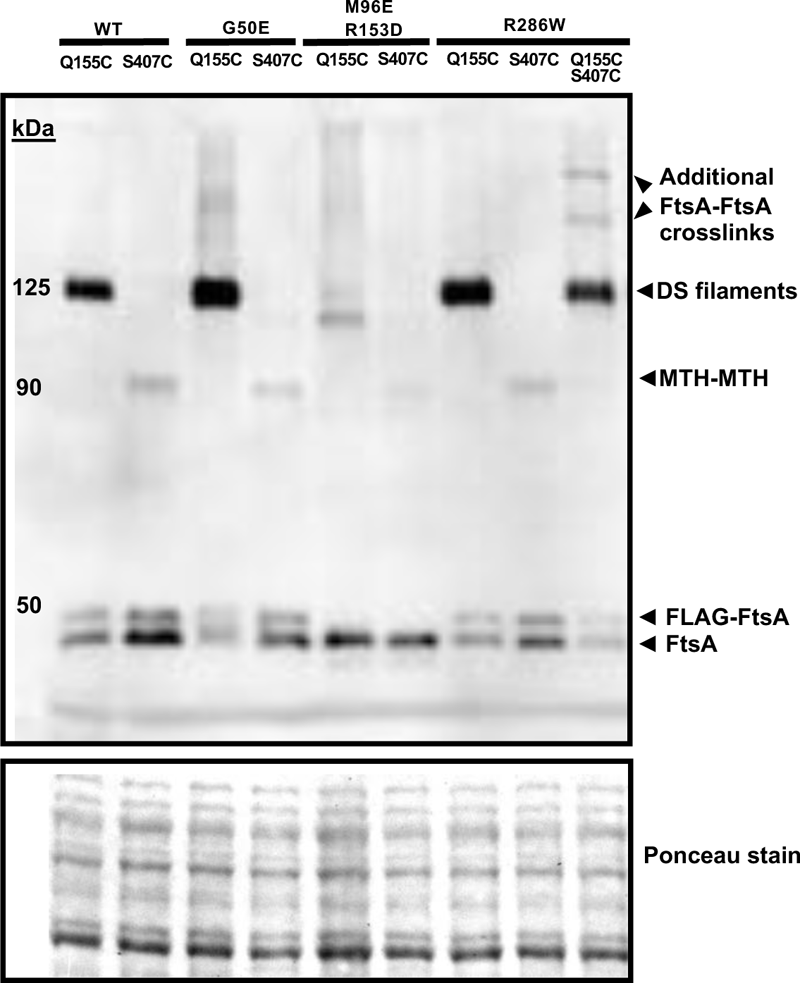
FtsA^M96E^ ^R153D^ is defective in forming DS filaments *in vivo*. Derivatives of plasmid pDSW210-FtsA^Q155C^ and/or FtsA^S407C^ in WM4952 carrying no additional residue changes (WT) or additional residue changes as shown, were assayed by *in vivo* cysteine crosslinking to identify DS filaments or MTH-MTH interactions, respectively. BMH was used as crosslinker. Crosslinked samples were separated by SDS-PAGE followed by immunoblotting with anti-FtsA. A portion of the Ponceau S-stained blot is shown below the immunoblot. All plasmid-expressed FtsA derivatives possess an N-terminal FLAG tag except for FtsA^M96E^ ^R153D^. The several larger crosslinked bands in the final lane suggest that FtsA protein carrying both Q155C S407C alterations can form both longitudinal and lateral crosslinks that result in higher-molecular weight structures.

We next wanted to test the effects of an FtsA variant reported to be defective in forming DS filaments. This variant of FtsA, FtsA^M96E^ ^R153D^, was previously designed based on the atomic structure to inhibit interaction between two antiparallel FtsA filaments (Fig. 2A) (30). In that study, a purified form of this variant formed robust minirings on lipid monolayers but, unlike WT FtsA, was unable to form typical DS filaments *in vitro* upon addition of purified FtsN_cyto_ (30).

To test the consequences of this defect on *E. coli* cell division, we first synthesized an internal portion of the *ftsA* gene encoding these two residue changes and replaced the WT *ftsA* in pSEB440, a medium copy plasmid with a *trc* promoter and relatively weak expression upon IPTG induction (47, 48), with the double mutant insert. This plasmid was chosen because of concerns that the double mutant might be dominant negative (see below). The double mutant *ftsA^M96E^ ^R153D^* was also cloned into the higher-copy number plasmid pDSW210 so that it could be expressed at higher levels from its P_Trc_ promoter for the cysteine crosslinking assays. Mutations encoding the Q155C and S407C residue changes were incorporated into the pDSW210-FtsA^M96E^ ^R153D^ plasmid to create triple mutants for cysteine crosslinking *in vivo*. As we suspected, cells carrying the pDSW210 derivatives of the double and triple mutants needed to be grown on 0.2 % glucose to repress the P_Trc_ promoter, as the cells grew poorly otherwise, likely due to a dominant negative effect of the double mutant (see below, and Fig. 5A).

**Fig. 5.**
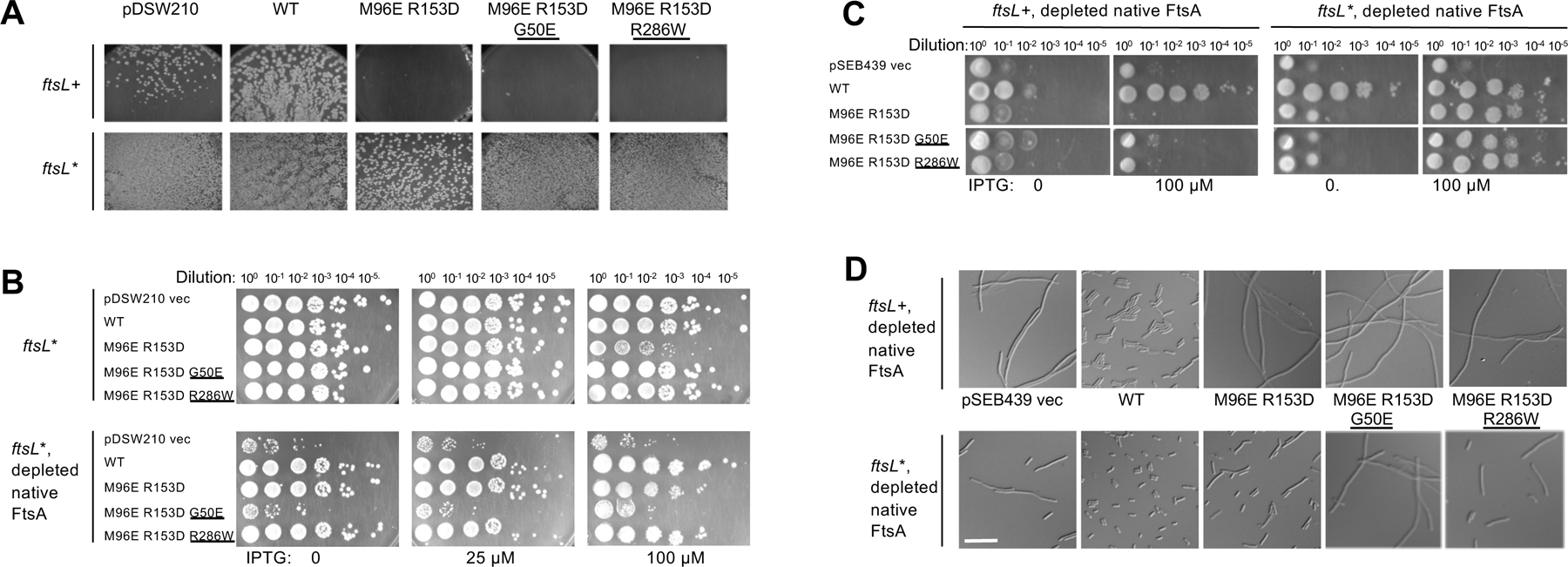
FtsA^M96E^ ^R153D^ is dominant negative and is unable to complement native FtsA except in the presence of the *ftsL*\* allele. *A*) Strains WM4952 (*ftsL*+, top row) or WM4953 (*ftsL*\*, bottom row) were transformed with pDSW210 vector or pDSW210 expressing WT or mutant FtsA derivatives (top), with no IPTG added, to maintain basal levels of expression. Images were taken after overnight incubation at 37°C. No transformants expressing FtsA^M96E^ ^R153D^ were obtained, even at uninduced levels of IPTG, unless the *ftsL*\* allele was present in the chromosome (WM4953) instead of *ftsL*+ (WM4952). The presence of G50E or R286W superfission alleles in *cis* did not permit viability without *ftsL*\*. B) Dominant negative effects of strains with *ftsL*\* and expressing WT FtsA or the mutant combinations from pDSW210 were tested at several different IPTG induction levels, with normal levels of native FtsA or depleted for native FtsA. In *ftsL*\* cells with normal levels of native FtsA (upper images), the dominant negative phenotype of FtsA^M96E^ ^R153D^, already strongly suppressed by *ftsL*\* as shown in (B), was further suppressed by R286W or G50E (detectable at 100 µM IPTG). When FtsA was depleted, FtsA^M96E^ ^R153D^ and FtsA^M96E^ ^R153D^ ^R286W^ strains were viable. However, when forced to be the sole functional FtsA, FtsA^M96E^ ^R153D^ ^G50E^ did not support viability, suggesting that the G50E allele is synthetically sick when combined with *ftsL*.* C) Controlled expression of FtsA and variants from pSEB439/pSEB440 after depletion of native FtsA in WM4952. WM4952 (*ftsL*+) or WM4953 (*ftsL*\*) cells carrying pSEB439 vector or expressing FtsA (pSEB440) or FtsA variants (pWM7215, pWM7496, pWM7522) were diluted and spotted onto agar plates and incubated at 30°C to deplete native FtsA, with either no IPTG or 100 µM IPTG to induce expression of the FtsA variants from the plasmid. D) Top row: Micrographs of overnight cultures of WM4952 at 30°C carrying pSEB440 derivatives as noted (+ 55 µM IPTG added to induce plasmid-expression of FtsA derivatives), indicating that FtsA^M96E^ ^R153D^ cannot substitute for native FtsA. Bottom row: same as top row except the strain background is the isogenic *ftsL*\* strain WM4953, which allows FtsA^M96E^ ^R153D^ and its derivatives to divide, although not completely normally. Scale bar, 10 µm.

Cysteine crosslinking experiments with FtsA^M96E^ ^R153D^ ^Q155C^ and FtsA^M96E^ ^R153D^ ^S407C^ derivatives showed that the ∼125 kDa band that represents crosslinking between Q155C residues in DS filaments was either undetectable or weak in repeat experiments (Fig. 4 and data not shown), indicating that the M96E and R153D residue changes strongly block formation of DS filaments *in vivo*. Crosslinking at S407C in the FtsA^M96E^ ^R153D^ double variant was also weaker compared to WT FtsA or FtsA^G50E^, but still detectable (Fig. 4 and data not shown). We conclude that the FtsA^M96E^ ^R153D^ double variant largely fails to assemble DS filaments *in vivo*, even when native FtsA is present.

### The dominant negative phenotype of FtsA^M96E^ ^R153D^ is suppressed by superfission alleles

As mentioned above, FtsA^M96E^ ^R153D^ derivatives were dominant negative, as certain strains could not be transformed by pDSW210-FtsA^M96E^ ^R153D^ in the absence of 0.2% glucose (Fig. 5A). This dominant negative phenotype could be explained if some defective subunits interacting with WT FtsA through longitudinal interactions prevented their co-assembly into DS filaments.

However, we hypothesized that perhaps a superfission allele such as *ftsL*\*, might suppress the dominant negative phenotype by stimulating septum synthesis, as it can rescue other cases of divisome inhibition (21, 22). Consistent with this idea, we found that pDSW210-FtsA^M96E^ ^R153D^ could efficiently be transformed into WM4953, which expresses *ftsL*\* from its native chromosomal locus and is isogenic with the *ftsL*+ WM4952 strain (Fig. 5A).

We surmised that another potential way to suppress the dominant negative effect of the double mutant was to incorporate G50E or R286W residue changes into the pDSW210-FtsA^M96E^ ^R153D^ plasmid to make two new triple variants. However, despite the known capability of G50E and R286W to suppress divisome defects, including suppression of other dominant negative mutants (19, 48, 49), they were unable to suppress the M96E R153D alterations that block DS filament formation. Instead, the triple variants remained dominant negative and failed to transform WM4952 (Fig. 5A) when selected on LB agar without added glucose to repress expression and under conditions where native FtsA was present (see below for additional description of this strain).

The inability of the G50E alteration to suppress the FtsA^M96E^ ^R153D^ dominant negative effect is consistent with the idea that the G50E residue change, despite its ability to promote DS filament formation in otherwise WT FtsA, does not have the capability to override the block to DS filament formation caused by FtsA^M96E^ ^R153D^. Moreover, the inability of R286W to suppress the FtsA^M96E^ ^R153D^ dominant negative phenotype suggests that FtsA needs to form DS filaments in order to complete the stimulation conferred by the R286W allele. This contrasts with the ability of the R286W residue change to suppress dominant negative alleles of FtsA such as FtsA^M71A^ (49) probably because the latter may favor an oligomeric state that occurs earlier in the process of cell division, such as divisome assembly, that is counteracted by R286W. In support of this idea, FtsA^M71A^ forms DS filaments significantly less well than WT FtsA (∼35% of WT levels) as measured by *in vivo* crosslinking at Q155C, but introduction of the R286W residue change to FtsA^M71A^ restores DS filament formation back to WT levels (Fig. S3).

As with the double variant, the triple variant constructs could be efficiently transformed into the *ftsL*\* strain WM4953, allowing us to investigate their effects on cell division and viability in more detail. We first asked whether FtsA^M96E^ ^R153D^ retained any dominant negative effect in the presence of *ftsL*\* compared with WT FtsA and the triple mutants. As seen in row 3 of the top panels in Fig. 5B, the double mutant retained some dominant negative properties compared with WT FtsA, which displayed no dominant negative effects at this induction level (row 2).

In contrast with the results from the *ftsL*+ WM4952 strain, the G50E or R286W residue changes in the FtsA^M96E^ ^R153D^ constructs in the *ftsL*\* WM4953 strain suppressed the dominant negative effects when native FtsA was present (Fig. 5B top panels, rows 4-5). This suppression by G50E or R286W was more pronounced than that achieved by *ftsL*\* alone, as shown by the normal viability of both triple mutants at 100 µM IPTG. This suggests that *ftsL*\* synergizes with G50E or R286W to compensate for the lack of FtsA DS filaments.

### The FtsA^M96E^ ^R153D^ variant is unable to replace FtsA unless rescued by *ftsL*\*

We then asked whether these FtsA variants were able to support cell division after native FtsA is depleted. Strains WM4952 and WM4953 harbor an *ftsA* null (*ftsA°*, frameshift) allele in the chromosome and a low-copy plasmid that expresses WT *ftsA* from the bacteriophage lambda P_R_ promoter along with its thermosensitive cI857 repressor. At temperatures >37°C, the repressor is thermo-inactivated, allowing WT FtsA to complement and cells to divide normally. However, shifting the temperature to 30°C or lower allows the repressor to inhibit P_R_, which severely depletes FtsA from cells (21), resulting in complete inhibition of cell division in the *ftsL*+ strain WM4952.

Previously it was shown that in WM4953 expressing the *ftsL*\* allele, when cellular FtsA is significantly depleted by growth at 30°C for several mass doublings, cell division is only mildly impaired (21). This reduced requirement for FtsA may occur because divisome activation by *ftsL*\* bypasses the requirement of most FtsA in the cell. Nonetheless, *ftsL*\* cannot bypass an *ftsA*° allele, indicating that some FtsA is still required for division in *ftsL*\* mutant cells (21), probably because FtsA is needed at least for recruitment of FtsL to the divisome. Moreover, the original *ftsL*\* allele (FtsL^E88K^) cannot bypass the loss of ZipA either (data not shown), presumably because ZipA is needed for recruitment of all downstream divisome proteins including FtsL.

To assess the functionality of FtsA variants, we first expressed them from pDSW210 derivatives under conditions (growth at 30°C) that depleted FtsA in either WM4952 or WM4953. As expected, cells with the pDSW210 vector control lost viability but cells expressing WT FtsA from pDSW210 retained normal viability (Fig. 5B, bottom panel, rows 1-2), even with no IPTG. This was expected, as there is sufficient leaky expression from pDSW210 with no IPTG to complement an *ftsA* null allele even in an *ftsL*+ strain (47). Notably, however, FtsA-depleted *ftsL*\* cells expressing FtsA^M96E^ ^R153D^ also exhibited normal viability with 0 or 25 µM IPTG, and showed a slight dominant negative effect at 100 µM IPTG (Fig. 5B, bottom panels, row 3), similar to the modest dominant negative effect in FtsA-replete cells mentioned above (Fig. 5B, top panels, row 3). This suggests that the double mutant retains at least partial function, provided that *ftsL*\* is present. Importantly, crosslinking experiments with the same FtsA^M96E^ ^R153D^ variant expressed from pDSW210 showed that it remains deficient in forming DS filaments in WM4953 (Fig. S2), ruling out the possibility that *ftsL*\* somehow allows restoration of DS filament formation.

The FtsA^M96E^ ^R153D^ ^R286W^ triple variant behaved similarly to the double variant, except there was no dominant negative effect (Fig. 5B, bottom panels, row 5). This suggests that when it is the main FtsA in the cell, the R286W variant synergizes with *ftsL*\* to bypass the block to DS filament formation. We were surprised, however, to find that the FtsA^M96E^ ^R153D^ ^G50E^ triple variant grew poorly at several IPTG induction levels after FtsA depletion (Fig. 5B, bottom panels, row 4). This finding suggests that when it is in the predominant FtsA in cells harboring the *ftsL*\* allele, the G50E residue change confers a dominant negative effect.

We then assessed the functionality of the FtsA variants when expressed from pSEB440, which allowed for finer control of *ftsA* expression. We used IPTG to induce expression of the different pSEB440-FtsA constructs under conditions (30°C growth) that depleted FtsA. As expected, depletion of native FtsA in *ftsL*+ WM4952 derivatives with the pSEB439 empty vector resulted in no viability (Fig. 5C, row 1) and long filamentous cells (Fig. 5D, top panels). WT FtsA restored full viability, but only after induction with IPTG (Fig. 5C row 2) and cell morphology was normal upon IPTG induction (Fig. 5D, top panels). Notably, although FtsA^M96E^ ^R153D^ and its triple variant derivatives could be introduced into WM4952 because of lower expression levels, they were unable to complement FtsA-depleted WM4952 even after IPTG induction (Fig. 5C, rows 3-5), and formed long filamentous cells (Fig. 5C left panels, rows 3-5; Fig. 5D, top panels). These results indicated that neither FtsA^M96E^ ^R153D^ nor the addition of G50E or R286W changes can replace native FtsA for function.

In contrast, in the *ftsL*\* WM4953 strain background, expression of FtsA^M96E^ ^R153D^ or FtsA^M96E^ ^R153D^ ^R286W^ after depletion of native FtsA resulted in normal viability on spot dilution plates (Fig. 5C right panels, rows 2,3,5), although this viability required IPTG induction (Fig. 5C, right panels), unlike WT FtsA. FtsA^M96E^ ^R153D^ ^G50E^ was slightly less viable (Fig. 5C right panels, row 4), consistent with the negative synergy with *ftsL*\* observed in Fig. 5B.

To confirm that these detrimental effects on viability were due to cell division defects, we compared the morphology of cells expressing FtsA^M96E^ ^R153D^ or the triple variants to that of cells expressing WT FtsA. Cells expressing FtsA^M96E^ ^R153D^ or FtsA^M96E^ ^R153D^ ^R286W^ divided, but less well than those expressing WT FtsA, with about half of the cells showing some degree of elongation (Fig. 5D, bottom panels 2, 3, 5). As predicted from the lower viability, cells expressing FtsA^M96E^ ^R153D^ ^G50E^ (Fig. 5D, fourth bottom panel) formed longer filaments, consistent with the negative synergy between G50E and *ftsL*\*.

### The FtsA^M96E^ ^R153D^ variant can function as the sole FtsA in cells carrying the *ftsL*\* allele

As FtsA^M96E^ ^R153D^ could allow cells with the *ftsL*\* allele to divide fairly well when WT FtsA was depleted, we then asked if FtsA^M96E^ ^R153D^ could function as the sole FtsA in the cell with no WT FtsA present. Using a P1 phage lysate carrying a *leuA*::Tn*10* tetracycline resistance (tet^R^) marker, which is ∼50% linked to the *ftsA°* allele by cotransduction (Fig. 6A), we transduced a WT *ftsL*+ strain carrying pSEB440-FtsA^M96E^ ^R153D^ to tet^R^ in the presence of 25-50 µM IPTG. We found that 100% of tet^R^ transductants tested retained the WT *ftsA* allele, indicating that FtsA^M96E^ ^R153D^ cannot function as the sole FtsA in an otherwise WT cell (data not shown) and consistent with its inability to complement WM4952 under FtsA depletion conditions (Fig. 5C, D)

**Fig. 6.**
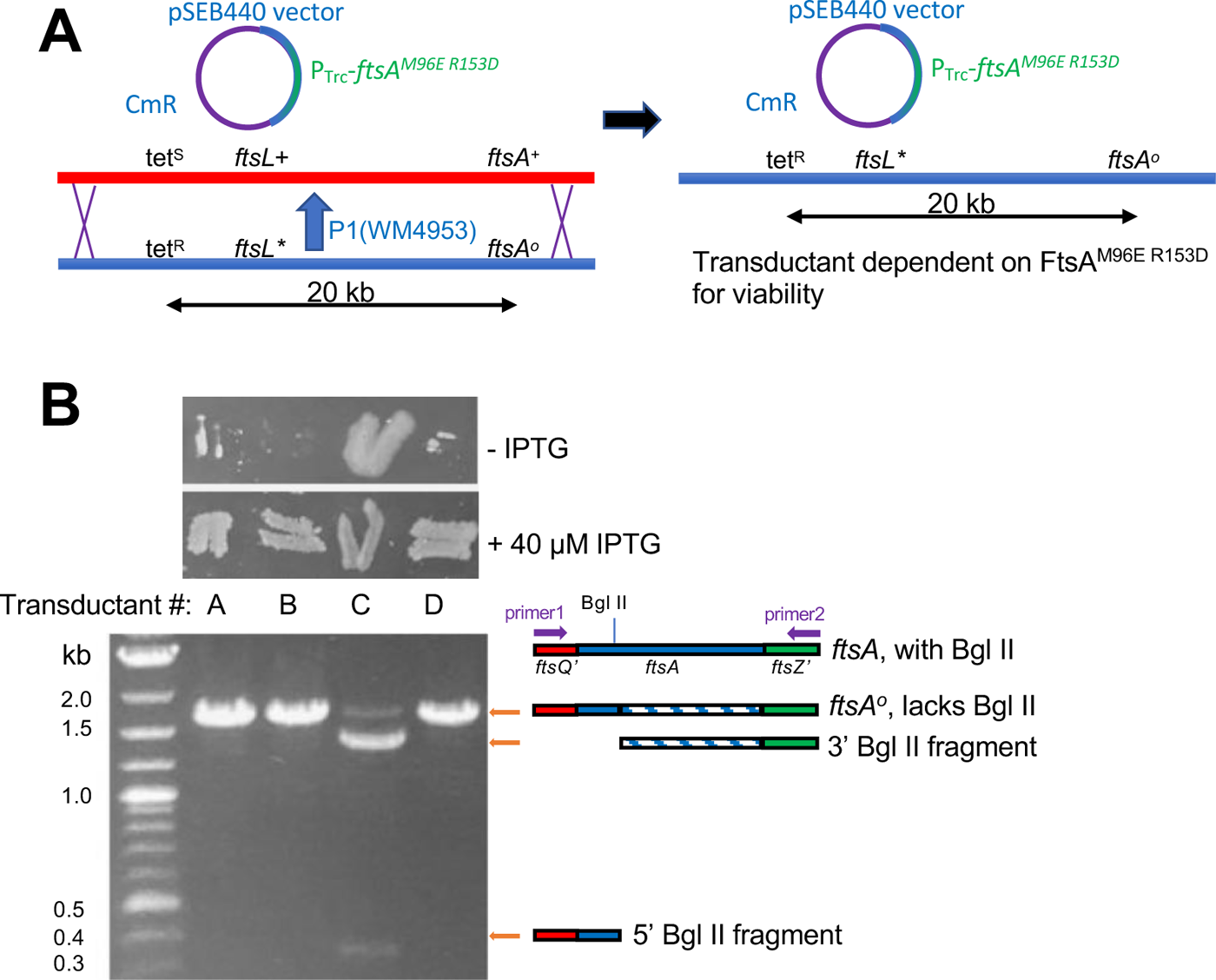
FtsA^M96E^ ^R153D^ can completely replace native FtsA in the cell when *ftsL*\* is present. (A) Scheme for P1 phage co-transduction of *ftsL*\* and *ftsA°* with the *tet^R^ leuO*::Tn*10* marker from WM4953 P1 lysate into a WT strain carrying pSEB440-FtsA^M96E^ ^R153D^ . The *ftsA°* and *ftsL*\* alleles are ∼90% and 50% linked to tetR, respectively. Double recombination necessary for cotransduction of *ftsA°* and *ftsL*\* alleles is denoted by X’s. The resulting Cm^R^ Tet^R^ transductant lacks WT *ftsA* and depends on FtsA^M96E^ ^R153D^ for viability. (B) Presence or absence of the *ftsA°*allele in four transductants from the experiment outlined in (A) confirmed by IPTG-dependence (patches of cells on LB agar ± IPTG, top) or Bgl II digested PCR products of chromosomal *ftsA* from transductant colonies (gel image, bottom). Isolates A, B and D have acquired *ftsA°*, and thus also must have acquired *ftsL*\* by cotransduction according to the scheme in (A).

We then repeated the transductions, this time with a phage lysate carrying *leuA*::Tn*10*, *ftsL*\* and *ftsA°* in that gene order (from WM4953), such that 100% of tet^R^ recipients receiving the *ftsA°* allele would also acquire the *ftsL*\* allele (Fig. 6A). With a WT strain carrying pSEB440-FtsA^M96E^ ^R153D^ as recipient, we reproducibly isolated ∼50% transductants that were dependent on IPTG for full viability, indicating that the cells now required FtsA^M96E^ ^R153D^ expressed from the plasmid for survival. A representative sample of 4 transductants out of dozens tested is shown in Fig. 6B. Microscopic examination of these cells revealed that 3 of the 4 were dependent on IPTG for cell division, and we confirmed that these isolates (A, B, D in Fig. 6B) had the *ftsA°* allele and that the plasmid in these strains retained the FtsA^M96E^ ^R153D^ residue changes (Fig. 6B and data not shown). The ∼50% cotransduction frequency of the *ftsA°* allele with the *leuA::Tn10* tet^R^ marker, as predicted from their 20 kb distance in the genome (Fig. 6A), also argues against the acquisition of extragenic suppressors, as these would be expected to arise at frequencies significantly lower than 50%.

### The FtsA^M96E^ ^R153D^ variant is capable of recruiting FtsN to the divisome

The ability of FtsL* to allow the FtsA^M96E^ ^R153D^ variant to function in cell division in the absence of WT FtsA strongly suggests that FtsA^M96E^ ^R153D^ can still successfully recruit downstream proteins, including FtsL*, to the divisome. Nonetheless, it is possible that the FtsA^M96E^ ^R153D^ variant, in the presence of *ftsL*\*, might be able to bypass some of the normal divisome assembly pathway. We explored this by testing if the late divisome protein FtsN could localize in *ftsA° ftsL*\* cells that were dependent on FtsA^M96E^ ^R153D^ for division and viability. We introduced pSEB440-FtsA^M96E^ ^R153D^ into a strain expressing a GFP-FtsN fusion at its native chromosomal locus (XTL970), then introduced the *ftsL* ftsA°* alleles into the chromosome by P1 transduction. The resulting strain (WM7518) was confirmed by whole genome sequencing to carry *ftsL*\* and *ftsA°* in the chromosome and to carry pSEB440-expressing FtsA^M96E^ ^R153D^ (data not shown). Complete viability of this strain was dependent on IPTG (Fig. 7A), as expected, although partial viability with no IPTG was also expected given the suppressing effects of *ftsL*\* on the amount of FtsA required for cell division as described above.

**Fig. 7.**
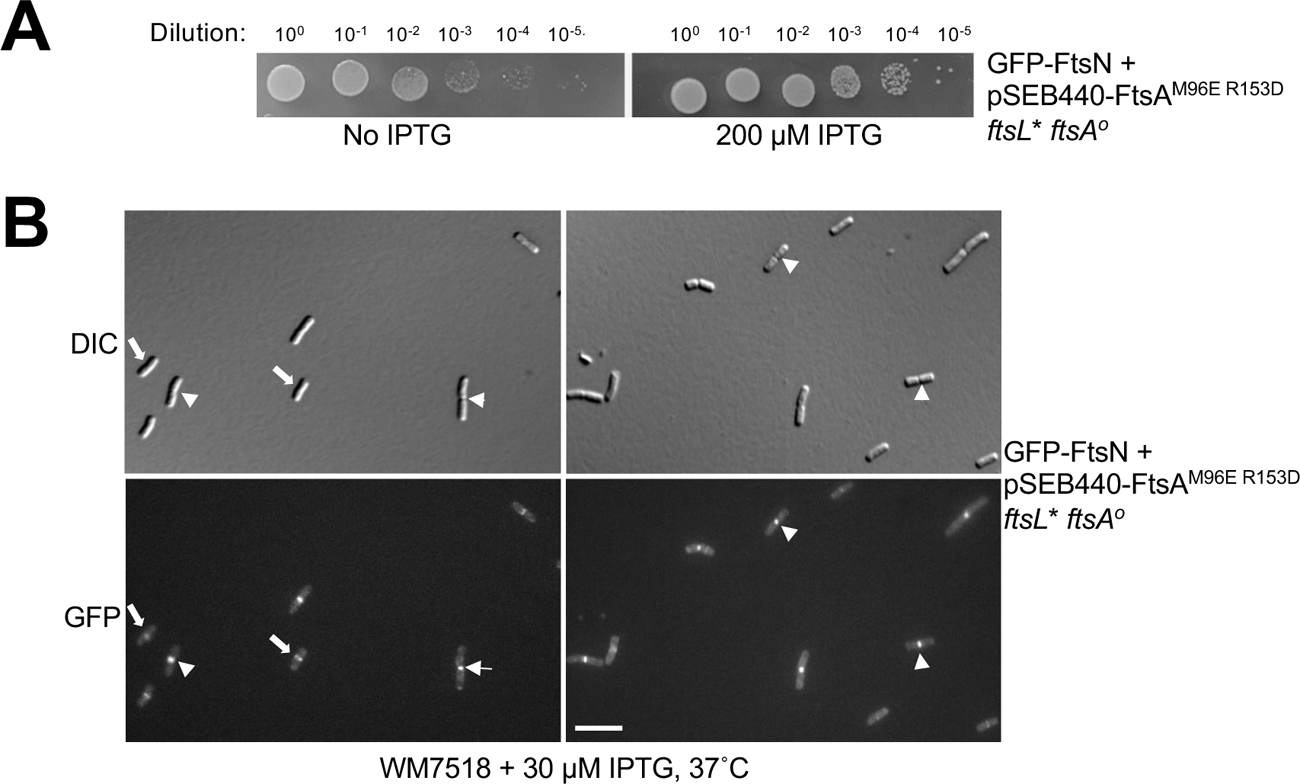
FtsA DS filaments are not required for divisome assembly or recruitment of FtsN. Strain WM7518 carries pSEB440-FtsA^M96E^ ^R153D^, expresses a GFP-FtsN fusion at the native *ftsN* chromosomal locus and harbors the *ftsL*\* and *ftsA°* alleles introduced by P1 transduction. (A) Confirmation that normal viability of WM7518 requires IPTG-dependent expression of FtsA^M96E^ ^R153D^, indicating that it is driving cell division in the absence of native FtsA. (B) The GFP-FtsN fusion localizes to nearly all divisomes in cells relying on FtsA^M96E^ ^R153D^ and *ftsL*\* for division. Two representative fields of cells (DIC and fluorescence) are shown, highlighting GFP-FtsN localization at early (arrows) or late (arrowheads) stages of septation. Scale bar, 5 µm.

Upon microscopic examination, we found robust GFP-FtsN fluorescent signals localized at divisomes in essentially all the cells (Fig. 7B), both early during the division process as well as in deeply constricting cells. This indicates that recruitment of later proteins by FtsA^M96E^ ^R153D^ is complete and suggests that FtsA-dependent recruitment of divisome proteins such as FtsN does not depend on the DS filament form of FtsA. The clear localization of GFP-FtsN to sharply constricting division septa also indicates that FtsN persists throughout cytokinesis, as is the case in WT cells (Soderstrom et al. 2016).

### The defect in forming DS filaments can be rescued by either *ftsL*\*\* or *ftsW*\* superfission alleles expressed from plasmids

To show independently that the *ftsL*\* superfission allele could rescue the FtsA^M96E^ ^R153D^ defect and be able to switch *ftsL*\* on when needed, we cloned a strong *ftsL* superfission allele, *ftsL^E88K^ ^G92D^* (22), which we call *ftsL**,* into the arabinose-inducible plasmid pBAD18. We then evaluated its capacity to restore cell division in the *ftsL*+ WM4952 strain carrying pSEB440-FtsA^M96E^ ^R153D^ after FtsA depletion at 30°C and induction with arabinose. A control strain expressing WT *ftsL* instead of *ftsL*\*\* from pBAD18 was also included for comparison.

We found that induction with IPTG and arabinose to express both FtsA^M96E^ ^R153D^ and FtsL** could fully rescue viability after FtsA depletion (Fig. 8A, bottom panels, row 2; Fig. 8B, left panel, row 3) and resulted in normal division of most FtsA-depleted cells in the population (Fig. 8C, left panel). As expected, IPTG induction of FtsA^M96E^ ^R153D^ alone did not help, and arabinose induction of FtsL** alone only partially suppressed FtsA depletion, consistent with previous observations that cells can divide with lower levels of FtsA when *ftsL*\* is present as mentioned above (21). In contrast to FtsL**, expression of WT FtsL with arabinose did not rescue viability of FtsA-depleted cells (Fig. 8A, bottom panels, row 2; Fig. 8C, left panel, row 2), and instead conferred weaker growth of FtsA-replete cells (Fig. 8A, top panels, row 2), indicating that excess WT FtsL is dominant negative. Overall, these data indicate that FtsA^M96E^ ^R153D^ can support cell division when FtsL** is co-produced.

**Fig. 8.**
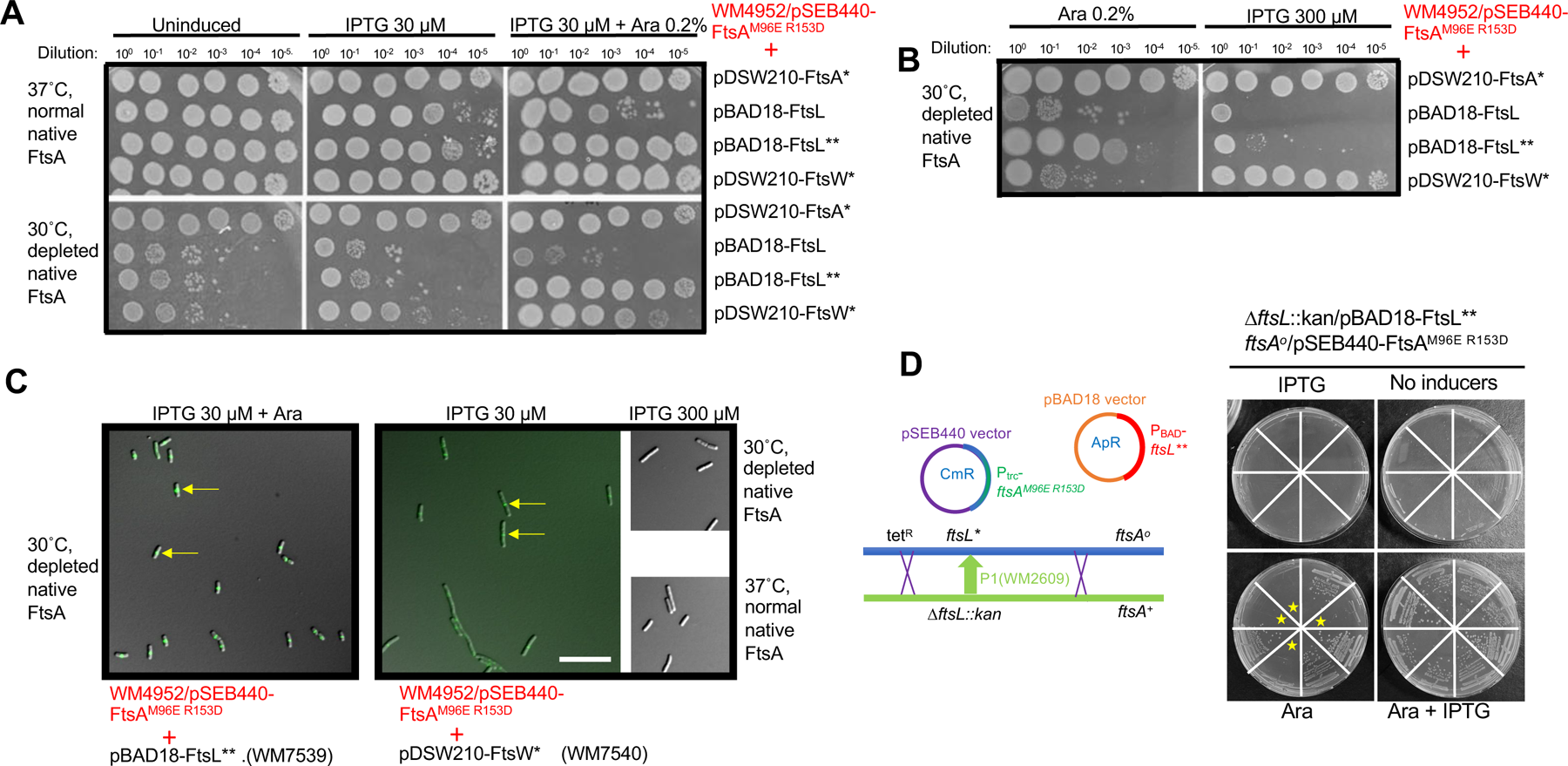
Superfission *ftsL*\*\* or *ftsW*\* alleles expressed from plasmids allow FtsA^M96E^ ^R153D^ to replace native FtsA. A) Plasmid pSEB440-FtsA^M96E^ ^R153D^ was introduced into the FtsA depletion strain WM4952 in combination with pDSW210-FtsA* (R286W, strain WM7599), pBAD18-FtsL (strain WM7600), pBAD18-FtsL** (E88K G92D, strain WM7539), or pDSW210-FtsW* (E289G, strain WM7540). Mid-logarithmic phase cultures grown at 37°C were serially diluted and spotted onto agar plates containing the inducers shown and incubated overnight at 37°C to maintain expression of the native FtsA, or 30°C to deplete native FtsA. IPTG is needed to express FtsA^M96E^ ^R153D^ or FtsW* from pSEB440 or pDSW210 plasmid, respectively. L-arabinose (Ara) is required to express FtsL or FtsL** from their plasmids. Full complementation by FtsA^M96E^ ^R153D^ after FtsA depletion was achieved with arabinose induction of FtsL** in combination with IPTG. There was sufficient leaky expression of FtsA* from pDSW210 to complement with no inducer. B) Induction of FtsL** with only arabinose partially rescued viability, but high level IPTG induction (300 µM) of FtsW* fully rescued viability. C) WM4952 derivatives with the plasmids indicated were grown in 30 µM IPTG + 0.2% arabinose at 30°C for 3.5 hours prior to imaging. Representative fields of cells are shown, after merging DIC and GFP channels. WM4952 expresses ZapA-GFP, a proxy for the Z ring (arrows), from its native locus. The short cell lengths for FtsL** and the mixture of short cells and moderate cell filamentation for FtsW* with 30 µM IPTG are consistent with the spot dilution results. Scale bar, 10 µm. D) FtsL** expressed from an arabinose-inducible promoter on a plasmid allows FtsA^M96E^ ^R153D^ to be the sole source of FtsA. A schematic for *ΔftsL::kan* transduction of WM7593 (*ftsL* ftsA°* pSEB440-FtsA^M96E^ ^R153D^ pBAD18-FtsL**) to replace the chromosomal *ftsL*\* allele with *ΔftsL::kan* is shown at the left; crossovers that would result in *ΔftsL::kan ftsA°* transductants (∼50% cotransduction frequency) are depicted. Eight transductants were picked at random and streaked on selective agar with or without IPTG (30 µM) or arabinose (0.2%) and incubated overnight at 30°C (shown at the right). All eight transductants were arabinose dependent; half (highlighted with stars) were IPTG dependent and thus depended on both FtsL** from pBAD18-FtsL** and FtsA from pSEB440-FtsA^M96E^ ^R153D^ for viability.

The ability of *ftsL*\* or *ftsL*\*\* to allow cell division independently of FtsA DS filaments suggested that DS filaments normally function to activate FtsWI-mediated septum synthesis--a role that can be bypassed if *ftsL\** activates FtsWI instead. We therefore reasoned that a *ftsW*\* superfission allele, which self-activates FtsWI, could also bypass the need for FtsA DS filaments. We cloned the strong *ftsW*\* allele *ftsW^E289G^* (Park et al. 2020) into the IPTG-inducible pDSW210 plasmid and assayed its ability to rescue viability of cells expressing FtsA^M96E^ ^R153D^ while depleted for native FtsA. Although low levels of IPTG (30 µM) were unable to rescue viability of colonies on plates (Fig. 8A), high levels of IPTG (300 µM) could restore full viability (Fig. 8B, C), suggesting that overexpression of *ftsW^E289G^*could bypass the need for FtsA DS filaments. These results indicate that both *ftsL\** and *ftsW\** superfission alleles can largely bypass the need for FtsA DS filaments, consistent with the idea that DS filaments are needed mainly for activation of FtsWI.

To confirm these results, we asked if FtsL** expressed from the pBAD18 plasmid could support viability of cells expressing FtsA^M96E^ ^R153D^ in the complete absence of WT FtsA. We transformed WM7518, which harbors the *ftsL*\* and *ftsA°* alleles in the chromosome and FtsA^M96E^ ^R153D^ expressed from pSEB440, with pBAD18-FtsL**, and then introduced the *ΔftsL::kan* knockout allele by P1 transduction, selecting on IPTG and arabinose to express FtsA^M96E^ ^R153D^ and FtsL**, respectively (Fig. 8D). All of the resulting transductants were arabinose-dependent as expected, but importantly, half of them were dependent on IPTG for viability and half were viable without IPTG (Fig. 8D), indicating that cell division depends on the expression of FtsA^M96E R153D^. (Half of the *ΔftsL::kan* transductants are expected to become IPTG-independent because the *ftsA*+ allele is ∼50% cotransducible with *ΔftsL::kan*). We obtained similar results when we used WM7317 instead of WM7518 (data not shown). Interestingly, we were repeatedly unable to introduce the *ftsA°* allele by P1 transduction into strains harboring WT *ftsL* and expressing both FtsL** and FtsA^M96E^ ^R153D^ from plasmids under induction conditions (data not shown). In summary, the viability of the *ΔftsL::kan* strain complemented by pBAD18-FtsL** suggests that (i) FtsL** produced from a plasmid can permit cell division in the absence of FtsA DS filaments and (ii) WT FtsL somehow interferes with *ftsL*\* superfission alleles, at least in the case of correcting FtsA DS filament defects.

### FtsA^G50E^ bypasses ZipA poorly compared with FtsA^R286W^

The FtsA^G50E^ allele, like other FtsA*-like variants, can complement an *ftsA°* allele and bypass ZipA (40). As ZipA is normally essential for recruiting downstream divisome proteins, potentially through its role in the disruption of FtsA minirings, this bypass of ZipA by FtsA^G50E^ is consistent with the latter being in a non-miniring oligomeric state. However, FtsA^G50E^ differs from FtsA^R286W^ in that the former seems to bypass the open arc form preferred by the latter variant. If the arc form is preferred for recruitment of FtsK and other downstream divisome proteins, we wondered if the constitutive DS filaments formed by FtsA^G50E^ might compromise its ability to bypass ZipA compared with FtsA^R286W^, particularly in a strain lacking any other source of FtsA.

To address this question, we used chromosomal *ftsA°* strains carrying pDSW210 derivatives expressing FtsA^R286W^, which should bypass ZipA efficiently; FtsA^S195P^, a thermosensitive variant which should fail to bypass ZipA even at the permissive temperature; and FtsA^G50E^, and transduced these with the *ΔzipA::kan* allele, selecting on plates containing 125 µM, 25 µM or 0 µM IPTG to assess the effects of different expression of the FtsA variants. As expected, FtsA^S195P^ could not be transduced with *ΔzipA::kan* under any induction condition (Fig. 9, bottom row), while FtsA^R286W^ transductants grew under all three conditions, but best with some IPTG induction (Fig. 9, top row). Interestingly, FtsA^G50E^ did not yield transductants without IPTG, and even with IPTG the transductants grew poorly compared to *ΔzipA::kan* mutants expressing FtsA^R286W^. Consistent with the observed colony phenotypes, cells of *ΔzipA::kan* transductants expressing FtsA^R286W^ and grown in 50 µM IPTG were mostly normal in length, whereas transductants expressing FtsA^G50E^ were uniformly filamentous (Fig. 9, right panels). Higher induction levels of IPTG did not enhance the ability of FtsA^G50E^ to bypass ZipA and instead inhibited growth (Fig. 9 and data not shown). These data suggest that when FtsA^G50E^ is the sole FtsA in the cell, it has a weakened ability to recruit downstream proteins compared with FtsA^R286W^.

**Fig. 9.**
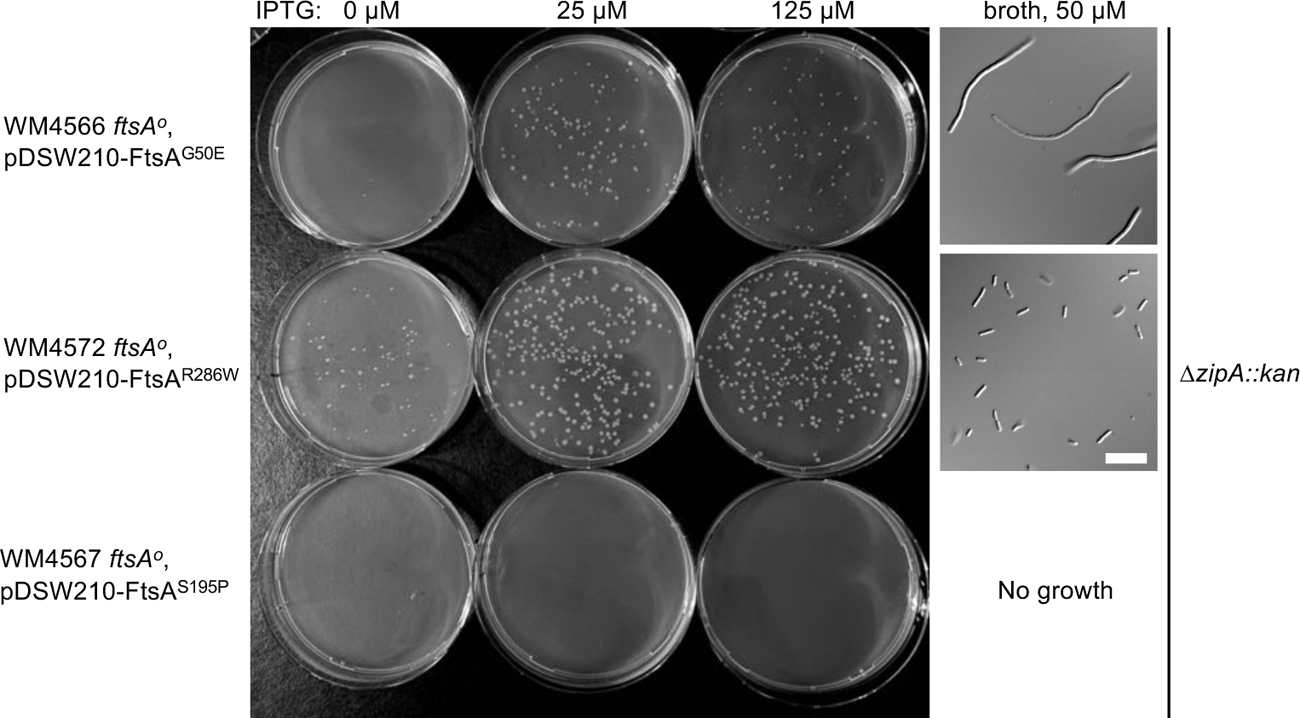
FtsA^G50E^ weakly bypasses ZipA compared with FtsA^R286W^. Strains WM4566, 4567 and 4572 were transduced with *ΔzipA::kan* P1 lysate and plated on LB Amp Kan plus concentrations of IPTG shown to express the various FtsAs from plasmids and incubated at 37°C overnight prior to imaging the agar plates. To examine cell morphology, one *ΔzipA::kan* transductant from FtsA^G50E^ or FtsA^R286W^ was inoculated in broth and grown in LB Amp Kan broth supplemented with 50 µM IPTG until reaching an OD_600_ of ∼0.5 (FtsA^R286W^) or 0.1 (FtsA^G50E^, which was concentrated ∼5x prior to viewing). Scale bar, 10 µm.

## DISCUSSION

Here we have characterized the role of DS filaments of FtsA in *E. coli* cell division. We provide evidence that DS filaments form late in the process of septation in *E. coli*, depend upon proper divisome assembly, and are important for divisome activation. The FtsA^M96E^ ^R153D^ variant, previously reported to be deficient in forming DS filaments *in vitro* in response to FtsN_cyto_, is unable to complement an *ftsA* null allele *in vivo* and is strongly dominant negative. This is consistent with the key role of DS filaments in FtsA’s cell division functions, and the dominant negative effect indicates that the loss of function of the FtsA^M96E^ ^R153D^ variant is not due to degradation, misfolding, or poor expression. Instead, we propose that the dominant negative effect occurs because assembly and stability of DS filaments require most FtsA subunits to interact at the lateral interface, and incorporation of even 1-2 subunits unable to form lateral interactions can destabilize the entire DS filament.

The most compelling evidence for the role of FtsA DS filaments in divisome activation comes from the ability of superfission alleles such as *ftsL*\* or *ftsW*\* to largely suppress the dominant negative effects of FtsA^M96E^ ^R153D^ and, even more strikingly, to allow it to function as the main source of FtsA in the cell. Superfission alleles prematurely promote septum synthesis by FtsWI, either by overriding normal signaling pathways from FtsQLB to FtsWI (*ftsL*\*) or by self-activation (*ftsW*\*) (12, 21–23). Therefore, as superfission alleles can bypass the need for FtsA DS filaments, it strongly suggests that DS filaments are normally involved in activating FtsWI. Although the molecular mechanisms underlying this activation are unclear, the known genetic interaction between FtsA and FtsW suggests that interaction between FtsA DS filaments and the cytoplasmic loops of FtsW, most likely loop 2 (50), triggers conformational changes in FtsW. It is attractive to postulate that these interactions in the cytoplasm, along with conformational changes in the periplasmic domains of FtsW promoted by FtsQLB, FtsI and FtsN, contribute synergistically to turn on FtsW’s glycosyltransferase activity (51–53).

Our results are consistent with the following model. FtsA initially forms minirings on the cytoplasmic membrane that represent a proto-ring checkpoint and are unable to recruit downstream divisome proteins, possibly because they lack free termini (Fig. 10). Then, responding to an unknown signal, ZipA and probably FtsX (33, 34) help to convert FtsA minirings into an “early” arc oligomeric state of FtsA, represented by FtsA^R286W^, which licenses recruitment of downstream divisome proteins. This model then postulates that the distinct “late” oligomeric state of FtsA is triggered by direct binding of FtsN_cyto_ to the FtsA 1C subdomain (30, 36, 37, 48), which tips the equilibrium towards DS filament assembly (Fig. 10). This conformational state is mimicked by FtsA variants such as FtsA^G50E^. In this model, DS filaments of FtsA are primed to activate FtsWI, likely by interacting more strongly with FtsW as described above; importantly, these DS filaments may have lost their ability to recruit divisome proteins for earlier stages of divisome assembly. The high levels of FtsN_cyto_ required to convert FtsA into DS filaments *in vitro* (30) suggests that other factors, including DS filaments themselves, may help to promote this transition *in vivo*.

**Fig. 10.**
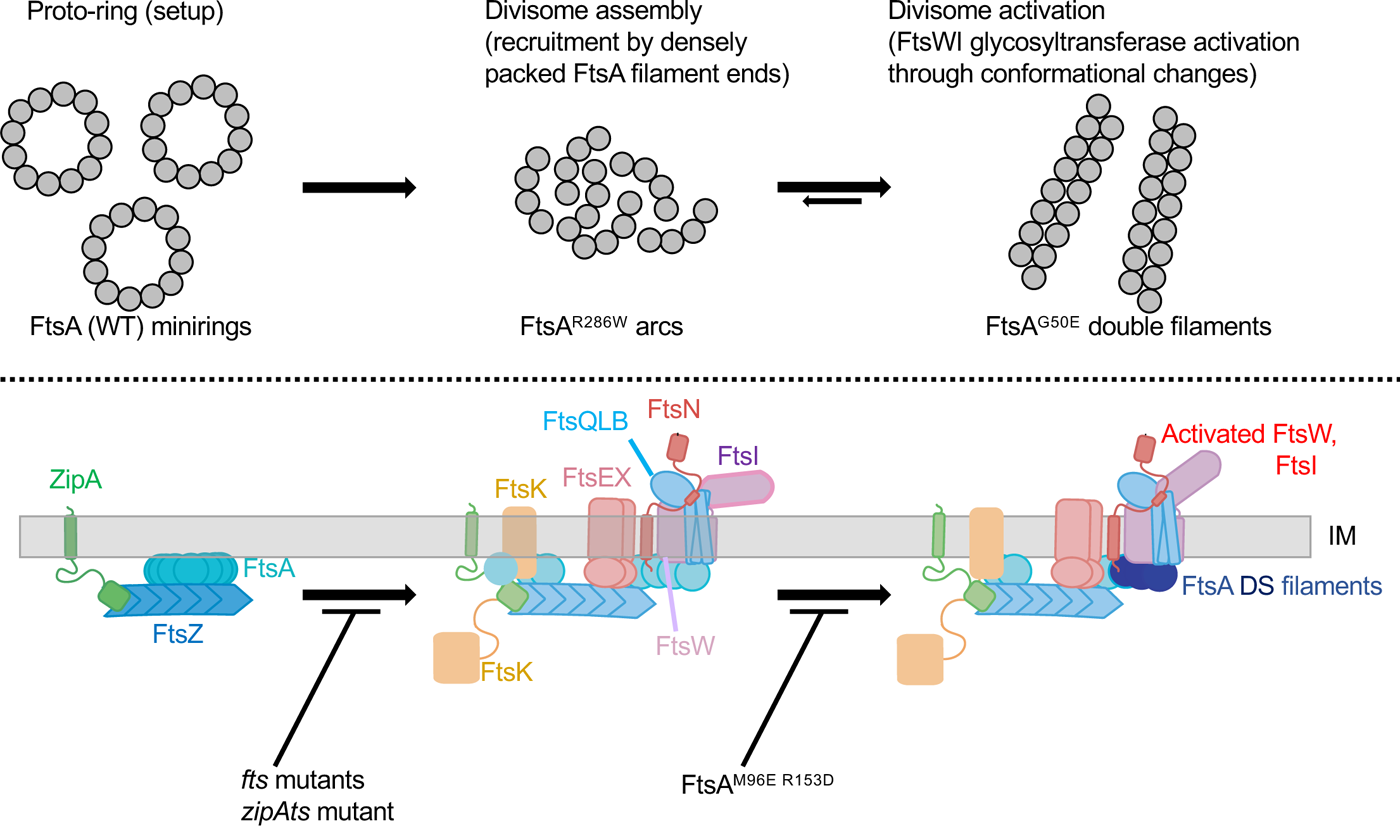
Proposed model of the effects of FtsA oligomeric states on divisome assembly and activation. FtsA minirings, arcs and DS filaments are shown above the proposed assembly/activation state of the divisome they affect, along with variants that favor the various FtsA oligomeric states. Note that there is no direct evidence yet for the existence of FtsA minirings and arcs *in vivo*. Mutants that block transitions between states are shown at the bottom.

The ability of FtsA^G50E^ to bypass ZipA weakly, as opposed to not at all like WT FtsA, suggests that the DS filaments are in equilibrium with arcs and that some arcs are present; this is consistent with the small percentage of non-DS filaments on lipid monolayers previously observed with purified FtsA^G50E^ (19, 30). Another possibility is that FtsA DS filaments can facilitate back-recruitment of divisome proteins, perhaps through FtsN (48, 54), but that this process is likely to be inefficient. The ability of FtsA^G50E^ to complement an *ftsA* null allele also suggests that ZipA can convert DS filaments into a divisome-recruiting form, or, perhaps more likely, that there are enough residual minirings (or other early structures) formed by FtsA^G50E^ that can be converted into recruiting forms by ZipA or other proteins. Interestingly, two other hypermorphic alleles of FtsA, FtsA^I143L^ and FtsA^E124A^, can largely bypass FtsN under certain conditions but are poor at bypassing ZipA (12, 16, 55). They share with FtsA^G50E^ a stronger tendency to form DS filaments on lipid membranes *in vitro*, as lower concentrations of FtsN_cyto_ are sufficient to convert minirings of FtsAI^I143L^ and FtsA^E124A^ to DS filaments compared with FtsA^R286W^ (30). Moreover, FtsA^Y139D^, which was previously isolated along with FtsA^G50E^ as a suppressor of FtsA^S195P^, forms a mixture of arcs and DS filaments, and like FtsA^G50E^ bypasses ZipA less efficiently than FtsA^R286W^ (19, 40). This is consistent with the idea that FtsA arcs bypass ZipA more readily than DS filaments. Despite their strong effects on FtsA oligomerization, the inability of the G50E and R286W residue changes to rescue the block to DS filaments caused by the M96E R153D lesions is consistent with the pathway shown in Fig. 10.

The negative synergy between G50E and *ftsL*\* is more difficult to explain, but may be related to the mechanism by which G50E forms mostly DS filaments. G50 is located at the longitudinal interface between FtsA subunits, and G50E therefore likely enhances lateral interactions through an allosteric effect (19). It is possible that this mechanism allows only a small number of FtsA arcs to be formed from minirings (see above) and thus perturbs recruitment of later divisome proteins. As FtsAG50E can complement an *ftsA°* allele, we speculate that a potential imbalance of divisome proteins might still manage to synthesize the division septum but would prone to failure if hyperstimulated by the *ftsL*\* allele. Clearly more studies of the costs of bypassing divisome checkpoints are needed.

A previous model for FtsA DS filaments proposed that this oligomeric form might play a role analogous to the DS filaments of MreB, which are postulated to act as rudders that sense and respond to membrane curvature, thus shepherding cell wall synthases to work in spatially restricted paths (28, 30, 39). In this analogy, DS filaments of FtsA would respond to the negative curvature of the invaginating septal membrane, thus spatially restricting FtsWI and perhaps FtsZ polymers to a narrow disc (30). This is an attractive model, particularly as it similar to the proposed mechanism of SepF, another FtsZ membrane anchor conserved in Gram positive bacteria and archaea, which can substitute for FtsA in *Bacillus subtilis* (5, 56, 57) and regulates septal morphology (58). The prevalence of DS filaments of FtsA late in the cell division process, observed in this study, is also consistent with the rudder model.

One of the corollaries of such a guiding mechanism for FtsA DS filaments is that they should be essential for septum formation. Indeed, the failure of FtsA^M96E^ ^R153D^ to complement an *ftsA* null allele indicates that DS filaments are essential. However, the surprising ability of *ftsL*\* or *ftsW*\* to bypass the requirement for DS filaments in cell division strongly suggests that their essential function is to activate FtsWI, and not as a crucial membrane guide for septum formation. It remains possible that other proteins have redundant roles with FtsA in guiding septum formation in *E. coli*. Given that *E. coli* septum formation involves a fast-moving treadmilling FtsZ track and a slower-moving septal synthase track comprising the FtsQLBWIN proteins (52, 59), and that FtsA can interact with proteins in both tracks (20, 35, 59) it is tempting to speculate that FtsA DS filaments may have a role in disengaging FtsQLBWIN from the FtsZ track. This would help to trigger septal synthase activity and help to spatially regulate it through an indirect mechanism.

## MATERIALS AND METHODS

### Strains, plasmids, and growth conditions

All strains and plasmids used for this study are listed in Table S1 in the supplemental material. Bacterial cultures were grown with Lennox lysogeny broth (LB) or agar plates unless otherwise indicated. Strains containing plasmids were supplemented as needed with ampicillin (50 µg/ml), chloramphenicol (15 µg/ml), tetracycline (10 µg/ml), kanamycin (25 µg/ml), IPTG (various concentrations) or L-arabinose (0.2%). Optical density of liquid cultures was assessed at 600 nm (OD_600_). Strains for plasmid construction were XL1-Blue, Top10, or WM7462 as detailed below.

### Construction of plasmids and strains

All primers are listed in Table S2. Cysteine substitutions at Q155C, S407C or D123C in *ftsA* were constructed by site-directed mutagenesis using primers 2564 + 2565, 2634 + 2635, or 2931 and 2932, respectively, on plasmid pWM2785 (pDSW210-FLAG-FtsA). G50E or R286W changes were made by site-directed mutagenesis using these derivatives as templates, with primers 2202 + 2203 for G50E and 2204 + 2205 for R286W.

The mutant gene encoding FtsA^M96E^ ^R153D^ was initially synthesized by Azenta Corp. as an internal fragment of the *ftsA* open reading frame, spanning nucleotides 217-496, and cloned into the pUC-GW-Kan plasmid vector. The internal *ftsA* gene fragment from this plasmid was the source for replacing the WT *ftsA* sequences in pSEB440 by Gibson assembly using pSEB440 vector primers 2659 and 2660, and internal primers 2661 and 2662, resulting in pSEB440-FtsA^M96E^ ^R153D^ (pWM7215).

To achieve higher expression levels, the *ftsA^M96E^ ^R153D^* mutant gene from pWM7215 was cloned into the high copy plasmid pDSW210 to make pWM7237. This was done by amplifying the *ftsA^M96E^ ^R153D^* from pWM7215 with primers 2580 + 2581, cleaving with *Kpn* I + *Asc* I, and cloning this fragment into plasmid pWM6246 (pDSW210-FtsA-msfGFP^sw^) (47), which was digested with *Kpn*I + *Asc*I (*Asc*I partial digestion, as there are two *Asc*I sites). The successfully cloned insert was screened for the replacement of a large portion of the *ftsA* gene and its msfGFP^sw^ insert with the smaller double mutant *ftsA* gene fragment lacking the msfGFP^sw^ insert and verified by sequencing.

To construct a derivative of *ftsA^M96E^ ^R153D^* for *in vivo* cysteine crosslinking, we introduced the Q155C change in the pDSW210-FLAG-FtsA derivative pWM7237 by site-directed mutagenesis using primers 2735 and 2736, or the S407C change using primers 2739 and 2740.

Other mutant combinations were made by site-directed mutagenesis. To introduce Q155C or S407C into FtsA^G50E^, plasmid pWM5733 (pDSW210-FLAG-FtsA^G50E^) was used as template for site-directed mutagenesis with primers 2564 and 2634, respectively. Sequences of pSEB440-FtsA derivatives were confirmed by sequencing the PCR product resulting from pSEB440-specific primers 2814 and 2815. The M71A variant was introduced into pWM7049 or pWM7239 by site-directed mutagenesis using primers 906 and 907.

To construct pBAD18-FtsL**, a DNA fragment containing the entire *E. coli ftsL* reading frame encoding the E88K and G92D residue changes, along with flanking sequences from upstream and downstream of the chromosomal *ftsL* gene including the *ftsL* translation initiation site, was synthesized by Twist Bioscience, Inc. This fragment was then cloned downstream of the P_BAD_ promoter on pBAD18-FtsL (pLD45) (60) by *in vivo* cloning in WM7462 (61), using primers 2892 and 2893 to amplify the *ftsL*\*\* gene and flanking sequences, and 2894 and 2895 to linearize the pBAD18 vector portion. To construct pDSW210-FtsW*, the entire *E. coli ftsW* open reading frame encoding the E289G residue change was synthesized by Twist Bioscience, Inc., and cloned into pDSW210 in frame with the ATG start codon in the vector by *in vivo* cloning using primers 2877 and 2878 to linearize pDSW210, and 2879 and 2880 as internal primers. The resulting plasmid expressed an N-terminally FLAG-tagged FtsW^E289G^ under P_Trc_ control.

The replacement of chromosomal *ftsL* with the kan^R^ cassette was achieved by recombineering, using the lambda Red-inducible strain DY329 (62) carrying pBAD18-FtsL (pLD45). This strain (WM2406) was electroporated with a PCR product amplified from a kan^R^ cassette and containing DNA flanking the *ftsL* gene using primers 712 + 713. Kan^R^ transformants were selected in the presence of arabinose and verified by their dependence on arabinose and by sequencing the junctions, and one strain was saved as WM2409. This strain was used to make a P1 phage lysate carrying the *ΔftsL::kan* allele that would be transduced into WM7593 to make WM7594.

The absence of WT *ftsA* after transduction with WM4953 lysate was initially verified in strains WM7317 and WM7518 by (i) the IPTG dependence of cell viability and (ii) amplification of the chromosomal *ftsQ’-ftsA-ftsZ*’ region from colony PCR confirming the lack of the unique Bgl II site within the N-terminus of chromosomal *ftsA* due to the frameshift mutation that marks the *ftsA°* allele (Fig. 6A). These two strains were also subject to whole genome sequencing (Plasmidsaurus, Inc.), which confirmed that they both harbor the *ftsL*\* and *ftsA°*alleles in their chromosomes and carried pSEB440 expressing FtsA^M96E^ ^R153D^. This allowed us to conclude that FtsA^M96E^ ^R153D^ can replace WT FtsA provided that *ftsL*\* is present.

To construct the *ftsA° ftsZ84*(ts) strain WM7412, WM1125 carrying pSEB440 was transduced with P1 lysate from WM4952 (*leuA*::Tn*10 ftsA°*), selecting for cm^R^ tet^R^ + IPTG. Because >95% of transductants receiving the *ftsA°* allele will also replace *ftsZ84*(ts) with *ftsZ*+, 100 transductants were screened for IPTG dependence and thermosensitivity. One transductant was isolated that was both IPTG dependent and thermosensitive, and confirmed to be *ftsA° ftsZ84*(ts).

The failure of *ftsL*\* to bypass ZipA was tested by transducing WM2991 (*zipA1ts*) with P1 lysate from WM4953 carrying *ftsL*\* closely linked to the *leuA::Tn10* marker and selecting for tet^R^ transductants at the permissive temperature of 30°C. Six transductants were then inoculated into LB broth + tetracycline and grown at 42°C for several hours, then examined microscopically. Cells in all six cultures were extremely filamentous. In addition, cells from the transductant colonies were longer than normal, a feature of the *zipA1* allele and in contrast to the very short cells typical of *ftsL*\* mutant strains.

Strains carrying pDSW210-FLAG-FtsA derivatives and a chromosomal *ftsA°*allele (WM4566, 4567, 4572) were grown at 30°C and tested for their ability to bypass *zipA* by transducing them with a *ΔzipA::kan* lysate derived from WM1657. Transductants were selected on LB + ampicillin + kanamycin at 30°C, supplemented with various concentrations of IPTG. Transductants of the WM4566 and WM4572 strains were examined microscopically by inoculating a single colony in corresponding liquid medium and growing until mid-logarithmic phase.

### Serial dilution plating assays

Overnight cultures were diluted 1:200 in fresh medium and grown for 2-3 h under permissive conditions to an OD_600_ of 0.2 to 0.4. Cultures were normalized by OD_600_ and then used to generate 10-fold serial dilutions in LB. Dilutions were spotted (5 µl) onto LB agar plates containing the indicated concentrations of IPTG or L-arabinose using a multichannel pipette. Plates were incubated overnight at temperatures indicated and then imaged.

### *In vivo* cysteine crosslinking

Overnight cultures were diluted 1:200 in fresh medium containing 50-100 µM IPTG (for expressing various cysteine-substituted *ftsA* genes from pDSW210) and grown to a final OD_600_ of 0.4 to 0.6 depending on the experiment. For the cell division synchronization experiment using the *ftsZ84* mutant strains carrying pDSW210-FLAG-FtsA, FtsA^Q155C^,FtsA^S407C^, or FtsA^D123C^, cultures were grown at 30°C until an OD_600_ of 0.2, at which point they were shifted to a 42°C shaking water bath for 45 min, then shifted back to 30°C. Aliquots from 0-, 10- and 20-min time points at 30°C were placed on ice and processed for crosslinking as described below. For the experiment with the *ftsA°* allele and *ftsQ1* and *zipA1* thermosensitive mutants, cultures were grown at 30°C until an OD_600_ of 0.2, at which point they were shifted to a 42°C shaking water bath for 45 min prior to harvesting for crosslinking. For the crosslinking experiment with WM4952 derivatives, cultures were all grown at 37°C.

Cultures were placed on ice then the equivalent of 1.0 OD_600_ of each was pelleted, washed with cold PBS, and resuspended in 50 µl cold PBS. BMB (1,4 bis-maleimidobutane) and BMH (bis-maleimidohexane) crosslinkers (ThermoFisher Inc.) or dBBR (dibromobimane) crosslinker (Sigma Aldrich) were prepared as 20 mM stock solutions in DMSO and added to each resuspended cell pellet to a final concentration of 0.5 mM. After 10 min of incubation, crosslinking was quenched by addition of 40 mM beta-mercaptoethanol (BME). Samples were pelleted and resuspended in 36.4 µl of freshly prepared resuspension buffer (50 mM Bis-Tris pH 7.0, 750 mM aminocaproic acid, 2 mg/ml lysozyme, 1 mM EDTA, 1.43 mM BME, 1x EDTA-free Pierce Protease Inhibitor [Thermo Scientific]) and incubated 20 min on ice. Dodecyl maltoside and SDS were each added to a final concentration of 0.9%, followed by an additional 20 min incubation on ice. Samples were then subjected to three cycles of 2-min incubation at 100°C followed by 2 min on dry ice, then pelleted in a microcentrifuge for 30 min. Supernatant fractions (40 µl) were combined with 5X sample buffer (312 mM Tris pH 6.8, 10% SDS, 50% glycerol, 25% BME) and boiled for 3 min prior to SDS-PAGE. Wells were generally loaded with 5 µl, or the equivalent of 0.1 OD_600_ of the original cell culture. Samples were stored at -20°C.

### Immunoblot analysis

Samples were boiled for 3 min prior to separation by SDS-polyacrylamide gel electrophoresis (PAGE). Samples were transferred to nitrocellulose membranes using a Mini Trans-Blot apparatus (Bio-Rad) at 200 mA for 55 min in Towbin transfer buffer containing 0.1% SDS. Blots were stained by Ponceau staining to assess total protein levels, then blocked using 3% bovine serum albumin (BSA) in Tris-buffered saline with Tween 20 (TBST) and incubated with 1:5000 rabbit anti-FtsA or mouse anti-FLAG (F1804, Sigma-Aldrich) antibodies. After washes, blots were incubated with 1:40,000 goat anti-rabbit-HRP or anti-mouse-HRP, developed with Bio-Rad Clarity ECL substrate and imaged using a Bio-Rad ChemiDoc MP system. The blots were also stained with Ponceau S (ThermoFisher) to visualize the levels of loaded protein samples across lanes. Band intensities were quantitated using ImageJ (63).

### Microscopy

Overnight cultures were generally diluted 1:200 in liquid media and grown to early or mid-logarithmic phase for imaging. Cells were either examined as wet mounts or mounted on agarose pads and imaged on an Olympus BX63 microscope equipped with a Hamamatsu C11440 ORCA-Spark digital complementary metal oxide semiconductor (CMOS) camera using cellSens software (Olympus). Images were analyzed using Fiji/ImageJ (63).

## Supporting information

Perkins et al. Supplemental material

## ACKNOWLEDGMENTS

This work was supported by NIH grant GM131705 to W.M. and funding from the MD Anderson UTHealth Graduate School of Biomedical Sciences to A.P. We thank Arindam Naha for initial cloning of the synthesized FtsA double mutant fragment into pSEB439 by Gibson assembly, Kara (Schoenemann) Hood and Brett Geissler for generating some of the strains and preliminary data, Xin-tian Li for providing us with the XTL970 strain, Joe Lutkenhaus and Sebastien Pichoff for providing us with the pSEB plasmids, Hironori Niki for providing the *in vivo* cloning strain, and Todd Cameron, Daniel Haeusser and Lorenzo Suigo for helpful discussions and advice.

