## Supplementary material for "Role of the antiparallel double-stranded filament form of FtsA in activating the *Escherichia coli* divisome": Perkins et al. Supplemental material

Contents:

Figures – Fig. S1-S3 with legends

Tables – Tables S1-S2

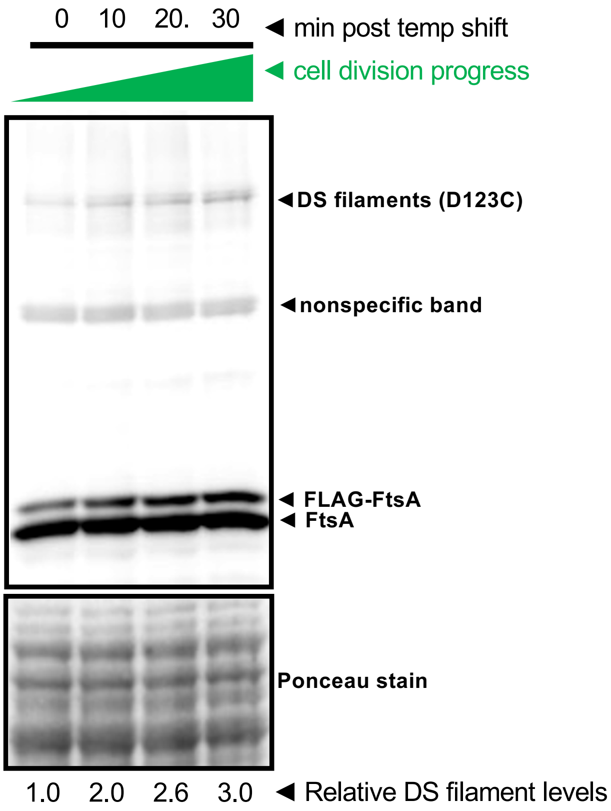

**Fig. S1. DS filaments measured by crosslinking at D123C increase later in cell division.** FtsA$^{D123C}$ in *ftsZ84*(ts) (WM7612) was used instead of FtsA$^{Q155C}$ to assay for DS filaments through *in vivo* crosslinking, probing immunoblots with anti-FtsA antibody, using the protocol described in Fig. 2B. As the distance between D123C residues in opposing antiparallel filaments is 3.7 Å (vs. 12.6 Å for Q155C), the shorter 5 Å dBBR crosslinker was used. A section of the same blot stained with Ponceau is shown below. Relative DS filament band intensities (ratio of crosslinked band to Ponceau stained protein at each time point, normalized to time zero) are shown at the bottom.

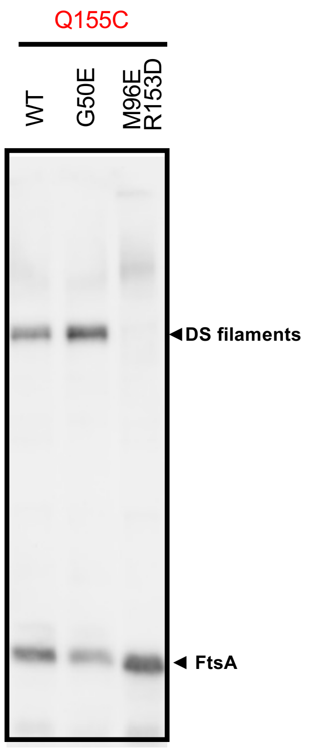

**Fig. S2. FtsA$^{M96E\ R153D}$ also fails to form DS filaments in an *ftsL** strain background.** WT, G50E or M96E R153D variants of FtsA also containing the Q155C residue for cysteine crosslinking (pWM7049, pWM7235, pWM7415) were each introduced into the *ftsL** WM4953 strain. After expression of each FtsA variant and crosslinking the Q155C residues with BMH, samples were separated by SDS-PAGE and probed with anti-FtsA on immunoblots.

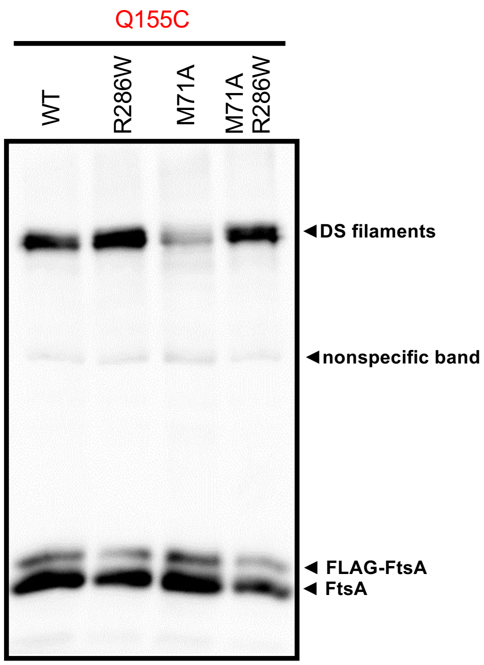

**Fig. S3. The previously characterized dominant negative variant FtsA$^{M71A}$ is deficient in forming DS filaments but can be rescued by R286W.** The FtsA variants expressed from pDSW210-FLAG shown (expressed from pWM7150, pWM7239, pWM7613, pWM7614, all in WM1074) were grown according to the standard crosslinking protocol and crosslinked with BMB prior to separation by SDS-PAGE and immunoblotting with anti-FtsA.

**Table S1.** Strains and plasmids used in this study.

| **Strain** | **Relevant genotype** | **Source or reference** |
|---|---|---|
| DY329 | W3110 Δ*lacU169 nadA::Tn10 gal490 pgl*Δ8 λcI*857*Δ*(cro-bioA)* | (1) |
| XTL970 | MG1655 *ΔlacI ΔaraE araFGH*<>spec, *lacY A177C*, *gfp-ftsN* | Xin-tian Li |
| Top10 | Cloning strain | Lab collection |
| XL1-Blue | Cloning strain | Lab collection |
| WM1074 | MG1655 *ΔlacU169* | Lab collection |
| WM1115 | *ftsA12 (ts)* in WM1074 | (2) |
| WM1125 | WM1074 *ftsZ84*(ts) | (3) |
| WM1657 | *ftsA*$^{R286W}$ Δ*zipA::aph* (Δ*zipA::kan)* | (2) |
| WM2406 | pBAD18-FtsL in DY329 recombineering strain | This study |
| WM2409 | WM2406 Δ*ftsL::kan* by recombineering; arabinose dependent | This study |
| WM2991 | W3110 *zipA1*(ts) | (4) |
| WM4566 | WM1074 *ftsA*$^{o}$ + pDSW210-FLAG-FtsA$^{G50E}$ | Lab collection |
| WM4567 | WM1074 *ftsA*$^{o}$ + pDSW210-FLAG-FtsA$^{S195P}$ | Lab collection |
| WM4572 | WM1074 *ftsA*$^{o}$ + pDSW210-FLAG-FtsA$^{R286W}$ | Lab collection |
| WM4661 | WM1074 *leuA::Tn10 ftsQ1*(ts) | (5) |
| WM4952 | MT78 (*leuA::*Tn*10 ftsA*$^{o}$, *zapA*-GFP, pSC101(ts)-P$_{R}$-*ftsA* | (6) |
| WM4953 | MT79 (MT78 with *ftsL** allele E88K) | (6) |
| WM7148 | pWM7049 (pDSW210FLAG-FtsA$^{Q155C}$) in *ftsQ1* (WM4661) | this study |
| WM7149 | pWM7049 (pDSW210FLAG-FtsA$^{Q155C}$) in *zipA1* (WM5337) | this study |
| WM7150 | WM1074 (pDSW210Flag-FtsA$^{Q155C}$) x P1(*ftsA*$^{o}$ *leuA::*Tn*10* | this study |
| WM7193 | WM1074 (pDSW210Flag-FtsA$^{S407C}$) x P1(*ftsA*$^{o}$ *leuA::*Tn*10* | this study |
| WM7237 | WM4953 + pDSW210-FtsA$^{M96E\ R153D}$ | this study |
| WM7305 | WM1074 + pSEB440-FtsA$^{M96E\ R153D}$ | this study |
| WM7317 | WM7305 x P1 (*leuA::*Tn*10 ftsL* ftsA*$^{o}$) | this study |
| WM7412 | WM1125 + pSEB440-FtsA x P1(*ftsA*$^{o}$ *leuA::*Tn*10*) | this study |
| WM7417 | WM4952 + pDSW210-FLAG-FtsA$^{G50E\ Q155C}$ | this study |
| WM7418 | WM4952 + pDSW210-FLAG-FtsA$^{G50E\ S407C}$ | this study |
| WM7419 | WM4952 + pDSW210-FtsA$^{M96E\ R153D\ Q155C}$ | this study |
| WM7420 | WM4952 + pDSW210-FtsA$^{M96E\ R153D\ S407C}$ | this study |
| WM7421 | WM4952+ pDSW210-FLAG-FtsA$^{R286W\ Q155C}$ | this study |
| WM7422 | WM4952 + pDSW210-FLAG-FtsA$^{R286W\ S407C}$ | this study |
| WM7423 | WM4952+ pDSW210-FLAG-FtsA$^{R286W\ Q155C\ S407C}$ | this study |
| WM7451 | WM4952 + pDSW210-FtsA$^{M96E\ R153D}$ | this study |
| WM7452 | WM4952 + pDSW210-FtsA$^{M96E\ R153D\ G50E}$ | this study |
| WM7453 | WM4952 + pDSW210-FtsA$^{M96E\ R153D\ R286W}$ | this study |

| | | |
|---|---|---|
| WM7454 | WM4953 + pDSW210-FtsA$^{\text{M96E R153D G50E}}$ | this study |
| WM7455 | WM4953 + pDSW210-FtsA$^{\text{M96E R153D R286W}}$ | this study |
| WM7462 | MG1655 *ΔhsdR ΔendA ΔrecA* (SN1187) *in vivo* cloning strain | (7) |
| WM7518 | pSEB440-FtsA$^{\text{M96E R153D}}$ in XTL970 x P1 (*leuA::*Tn*10 ftsL* ftsA*$^{o}$*)* | this study |
| WM7519 | pSEB440-FtsA$^{\text{R260W}}$ in XTL970 x P1 (*leuA::*Tn*10 ftsL* ftsA*$^{o}$*)* | this study |
| WM7522 | WM4952 + pSEB440-FtsA$^{\text{M96E R153D G50E}}$ | this study |
| WM7539 | WM4952 + pSEB440-FtsA$^{\text{M96E R153D}}$ + pBAD18-FtsL** | this study |
| WM7540 | WM4952 + pSEB440-FtsA$^{\text{M96E R153D}}$ + pDSW210-FtsW* | this study |
| WM7547 | WM1125 + pDSW210-FLAG-FtsA$^{\text{Q155C}}$ x P1(*ftsA*$^{o}$ *leuA::*Tn*10* | this study |
| WM7550 | WM4952 + pDSW210-FLAG-FtsA$^{\text{Q155C}}$ (pWM7049) | this study |
| WM7551 | WM4953 + pDSW210-FLAG-FtsA$^{\text{Q155C}}$ (pWM7049) | this study |
| WM7593 | WM7518 + pBAD18-FtsL** | this study |
| WM7594 | WM7593 ∆*ftsL::kan* (x P1 WM2409) | this study |
| WM7599 | WM4952 + pSEB440-FtsA$^{\text{M96E R153D}}$ + pDSW210-FtsA* | this study |
| WM7600 | WM4952 + pSEB440-FtsA$^{\text{M96E R153D}}$ + pBAD18-FtsL | this study |
| WM7604 | WM4953 + pSEB440 | this study |
| WM7605 | WM4953 + pSEB440- FtsA$^{\text{M96E R153D}}$ | this study |
| WM7606 | WM4953 + pSEB440- FtsA$^{\text{M96E R153D G50E}}$ | this study |
| WM7607 | WM4953 + pSEB440- FtsA$^{\text{M96E R153D R286W}}$ | this study |
| WM7612 | WM1125 + pDSW210-FLAG-FtsA$^{\text{D123C}}$ | this study |
| **Plasmid** | **Description** | **Source or reference** |
| pUC-GW-Kan | Cloning vector | Azenta Corp. |
| pDSW210 | *colE1* plasmid with weakened P*trc* promoter, Amp$^{R}$ | (8) |
| pKG110 | pACYC184 derivative containing the *nahG* promoter, Cm$^{R}$ | (5) |
| pSEB439 | pACYC184 derivative with a Ptrc promoter | (9) |
| pSEB440 | pSEB439 expressing *ftsA* | (9) |
| pWM2785 | pDSW210-FLAG-FtsA | (10) |
| pBAD18 | *colE1* plasmid with *araBAD* promoter | (11) |
| pBAD18-FtsL | pBAD18 expressing *ftsL*, arabinose control (pLD45) | (12) |
| pBAD18-FtsL** | pBAD18 expressing *ftsL*$^{\textit{E88K G92D}}$ | this study |
| pDSW210-FtsW* | pDSW210-FtsW$^{\text{E289G}}$ | this study |
| pWM5732 | pDSW210-FLAG-FtsA$^{\text{R286W}}$ | this study |
| pWM5733 | pDSW210-FLAG-FtsA$^{\text{G50E}}$ | this study |
| pWM6246 | pDSW210-FtsA-msfGFP$^{\text{sw}}$ | (13) |
| pWM7049 | pDSW210-FLAG-FtsA$^{\text{Q155C}}$ | this study |
| pWM7181 | pDSW210-FLAG-FtsA$^{\text{S407C}}$ | this study |
| pWM7159 | pUC-GW-Kan-*ftsA* internal fragment encoding M96E R153D | this study |
| pWM7215 | pSEB440- FtsA$^{\text{M96E R153D}}$ | this study |

| | | |
|---|---|---|
| pWM7235 | pDSW210-FLAG-FtsA$^{G50E\ Q155C}$ | this study |
| pWM7236 | pDSW210-FLAG-FtsA$^{G50E\ S407C}$ | this study |
| pWM7237 | pDSW210-FtsA$^{M96E\ R153D}$ | this study |
| pWM7239 | pDSW210-FLAG-FtsA$^{R286W\ Q155C}$ | this study |
| pWM7240 | pDSW210-FLAG-FtsA$^{R286W\ S407C}$ | this study |
| pWM7415 | pDSW210-FLAG-FtsA$^{M96E\ R153D\ Q155C}$ | this study |
| pWM7423 | pDSW210-FLAG-FtsA$^{Q155C\ R286W\ S407C}$ | this study |
| pWM7496 | pSEB440- FtsA$^{M96E\ R153D\ R286W}$ | this study |
| pWM7522 | pSEB440- FtsA$^{M96E\ R153D\ G50E}$ | this study |
| pWM7612 | pDSW210-FLAG-FtsA$^{D123C}$ | this study |
| pWM7613 | pDSW210-FLAG-FtsA$^{M71A\ Q155C}$ | this study |
| pWM7614 | pDSW210-FLAG-FtsA$^{M71A\ R286W\ Q155C}$ | this study |

**Table S2.** Primers used in this study.

| Primer # | 5'-3' nucleotide sequence |
|---|---|
| 712 ΔftsL-f | TGCAGAGAGGACGAATGCATGATCAGCAGAGTGACAGAAGGCCACGTTGTGTCTC |
| 713 ΔftsL-r | GCGTTTTCGCCGCTGCTTTCATGCGTCGCGTTTATCCTTAGAAAAACTCATCGAGC |
| 906 | CATTGACCAGGCAGAATTGGCCGCAGATTGTCAGATCTC |
| 907 | GAGATCTGACAATCTGCGGCCAATTCTGCCTGGTCAATG |
| 2202 | GGATAAAGGCGAAGTGAACGACCTCGAATC |
| 2203 | ATACCACGCGACGGGCAG |
| 2204 | TCGTCCGCCATGGAGTCTGCAAC |
| 2205 | CCACCTACGCTCGGCACTTC |
| 2564 | CGTGCGGATGTGTGCAAAAGTGCACC |
| 2565 | CCCGAAAGTCCTACCGGA |
| 2580 | GGTAGTAGGACTGGAGATTGGTACC |
| 2581 | CTCCTGAGCATAATCCGTTAAACC |
| 2634 | ATCAGTTGGCTGCTGGATCAAGCG |
| 2635 | GCTGTAACACGTTTTTCTAC |
| 2659 | ATATGGCGAAAAACATCGTCAA |
| 2660 | TGCCATCAATTCTGCCTGGTC |
| 2661 | CATTGACCAGGCAGAATTGATGGCAGATTGTCAGATCTCTTCGG |
| 2662 | GCTTTGACGATGTTTTTCGCCATATCGTTGTGACATGTGATCAGG |
| 2735 | GACTTTCGGGCGTCGATATGTGTGCAAAAGTGCACCTGATCAC |
| 2736 | GTGATCAGGTGCACTTTTGCACACATATCGACGCCCGAAAGTC |
| 2739 | GTTACAGCATCAGTTGGCTGCTGGATCAAGCGACTC |
| 2740 | GAGTCGCTTGATCCAGCAGCCAACTGATGCTGTAAC |
| 2814 | TTTTTATGAGGCCGCATGCA |
| 2815 | TGTTGTTCTGCCTGTGCCTATTCCA |
| 2877 | GGCTGTTTTGGCGGATGAGAGAAG |
| 2878 | GGCTGTTTTGGCGGATGAGAGAAG |
| 2879 | GGATCCTCTAGAATGATCAAGGCGATGCGTTTATCTCTCCCTCGCCTG |
| 2880 | CTTCTCTCATCCGCCAAAACAGCCCTGCAGTCATCGTGAACCTCGTAC |
| 2892 | CCATACCCGTTTTTTTGGGCTAGCGAATTCCGTATTGCAGAGAGGACGAATG |
| 2893 | CATCCGCCAAAACAGCCAAGCTTGCATACTCTAGAGCATGCGTCGCGTTTATCC |
| 2894 | ATGCAAGCTTGGCTGTTTTGGCGGATG |
| 2895 | GAATTCGCTAGCCCAAAAAAACGGGTATGG |
| 2931 | GCGAAATCGGTGCGTGTGCGCTGTGAGCATCGTGTGCTGCATGTG |
| 2932 | CACATGCAGCACACGATGCTCACAGCGCACACGCACCGATTTCGC |
